# Myosin A and F-Actin play a critical role in mitochondrial dynamics and inheritance in *Toxoplasma gondii*

**DOI:** 10.1101/2024.03.18.585462

**Authors:** Rodolpho Ornitz Oliveira Souza, Chunlin Yang, Gustavo Arrizabalaga

**Author notes:** Corresponding author Address: 635 Barnhill Drive, MS Building, Room MS A418, Indianapolis, IN, 46202.

## Abstract

The single mitochondrion of the obligate intracellular parasite *Toxoplasma gondii* is highly dynamic. Toxoplasma’s mitochondrion changes morphology as the parasite moves from the intracellular to the extracellular environment and during division. Toxoplasma’s mitochondrial dynamic is dependent on an outer mitochondrion membrane-associated protein LMF1 and its interaction with IMC10, a protein localized at the inner membrane complex (IMC). In the absence of either LMF1 or IMC10, parasites have defective mitochondrial morphology and inheritance defects. As little is known about mitochondrial inheritance in *Toxoplasma*, we have used the LMF1/IMC10 tethering complex as an entry point to dissect the machinery behind this process. Using a yeast two-hybrid screen, we previously identified Myosin A (MyoA) as a putative interactor of LMF1. Although MyoA is known to be located at the parasite’s pellicle, we now show through ultrastructure expansion microscopy (U-ExM) that this protein accumulates around the mitochondrion in the late stages of parasite division. Parasites lacking MyoA show defective mitochondrial morphology and a delay in mitochondrion delivery to the daughter parasite buds during division, indicating that this protein is involved in organellar inheritance. Disruption of the parasite’s actin network also affects mitochondrion morphology. We also show that parasite-extracted mitochondrion vesicles interact with actin filaments. Interestingly, mitochondrion vesicles extracted out of parasites lacking LMF1 pulled down less actin, showing that LMF1 might be important for mitochondrion and actin interaction. Accordingly, we are showing for the first time that actin and Myosin A are important for *Toxoplasma* mitochondrial morphology and inheritance.

## INTRODUCTION

*Toxoplasma gondii* is an intracellular parasite that can infect any nucleated cell, and it is known to infect almost all homeothermic animals; humans among them. Besides its importance as an opportunistic pathogen in immunocompromised individuals, *Toxoplasma* possesses unique cell biology whose study can lead to a broader understanding of eukaryotic evolution and identify new targets for therapeutic interventions. *Toxoplasma* divides by a specialized process known as endodyogeny. In this process, two daughter cells are formed within a mother cell (1). During this process, the pellicle, which is made up of the parasite membrane, the inner membrane complex (IMC), and a network of intermediate filaments, plays an important role in the formation of new cells. Primarily, two new IMCs form within the mother parasite and provide the scaffold for the division and distribution of organelles and parasite structures (2). While several organelles, such as the rhoptries and micronemes, are synthesized *de novo,* many others are equally segregated between the two nascent daughter parasites (2). Of particular interest is the tubular mitochondrion of the parasite, which is essential for parasite survival and a validated drug target. Since the *Toxoplasma* mitochondrion is a single-copy organelle, its division and inheritance are tightly associated with the parasite cell cycle and division. In non-dividing parasites, the mitochondrion is in a so-called lasso shape and is distributed along the periphery of the parasite, where it establishes contact sites with the IMC (2, 3). Interestingly, the mitochondrion is the last organelle to divide during endodyogeny (2, 4). Late in division, the mitochondrion appears to dissociate from the mother parasite IMC and develops branches that extend along its length. These branches enter the daughter cells at the last point of division and quickly spread inside the new cells, reestablishing the lasso shape. Despite the importance of the mitochondrion to parasite survival, surprisingly, little is known about the proteins driving and regulating mitochondrial division and inheritance.

In recent years, progress has been made in identifying the proteins involved in maintaining the mitochondrion in a lasso shape at the periphery of the parasite. Our laboratory characterized a mitochondrion-associated protein named LMF1 (short for Lasso Maintenance Factor 1) that appears to mediate the contact site between the outer mitochondrial membrane and the IMC. In the absence of LMF1, the mitochondrion loses its lasso shape and collapses towards one of the ends of the parasite (5). To mediate the contact site, LMF1 interacts with IMC10, a protein located in the parasite’s IMC. Similarly to LMF1, loss of IMC10 causes mitochondrion morphological defects, in addition to a few division-related defects (6). Interestingly, loss of either LMF1 or IMC10 results in parasites that lack mitochondrion and in mitochondrial material outside of the parasites. Thus, it appears that the LMF1/IMC10 tether might be important not only for maintaining the lasso shape in non-dividing parasites but also for the distribution of the mitochondrion among the daughter parasites. Accordingly, we observe that IMC10 and LMF1 form a tether immediately upon mitochondrion entrance into the daughter cells (6). It is plausible that the formation of the tether is critical for positioning the mitochondrion for entry into the daughter cell using the IMC as a scaffold or anchor for that process. Nonetheless, what provides the force to pull or push the mitochondrion into the daughter cells is unknown.

In other eukaryotes, actomyosin plays an important role in mitosis and mitochondrion segregation (7). It has been shown that the actin-myosin system is important for mitochondrial fission (8), movements (9), and mtDNA maintenance (10) in mammalian cells. In yeast, mitochondrial inheritance depends on the actin-myosin network (11–13). Myosin 2 is a cargo-binding myosin responsible for mitochondrial movements into the newly formed cell during budding (14–17). Recent studies have shown the importance of unconventional myosins and actin in the segregation of the apicoplast, a non-photosynthetic plastid in *Toxoplasma* (18–20). Intriguingly, the yeast two-hybrid screen that identified IMC10 as an interactor of LMF1 also identified the unconventional motor MyoA as a putative LMF1 interactor. MyoA is an unconventional motor that belongs to Class XIV of myosins. It is a fast, single-headed, and non-processive motor that drives parasite gliding motility (21, 22). In this study, we investigate the role of MyoA as a motor protein involved in mitochondrion inheritance. Cells lacking MyoA are defective in mitochondrion positioning and distribution to the newly formed cells. Furthermore, we found that *Toxoplasma* mitochondrion interacts with F-actin in vitro, and its positioning relies on proper actin dynamics. Thus, our studies have revealed a novel role for MyoA in mitochondrial morphology and division, expanding our understanding of how *Toxoplasma* ensures the inheritance of its single mitochondrion during division.

## RESULTS

### Myosin A interacts with LMF1

Previous work used a Yeast Two-Hybrid (Y2H) interaction screen to identify LMF1 interactors (6) and obtained a list of 15 proteins, which were prioritized based on cellular localization. Among these proteins, three proteins were located within the pellicle, Myosin-A (MyoA) among them (Fig. 1A). Given the fact that MyoA is a motor protein, we were interested in investigating the possible role of this protein in the parasite’s mitochondrial dynamics. Immunoprecipitation (IP) of LMF1-HA from intracellular parasites using anti-HA magnetic beads followed by mass spectrometry analysis, revealed peptides corresponding to MyoA as well as to the myosin light chain MLC1, providing further evidence of an interaction between LMF1 and MyoA (Fig. 1B). Interestingly, this interaction could not be detected when the IP was performed using extracellular parasites, as no peptides could be detected for MyoA or MLC1.

**Figure 1.**
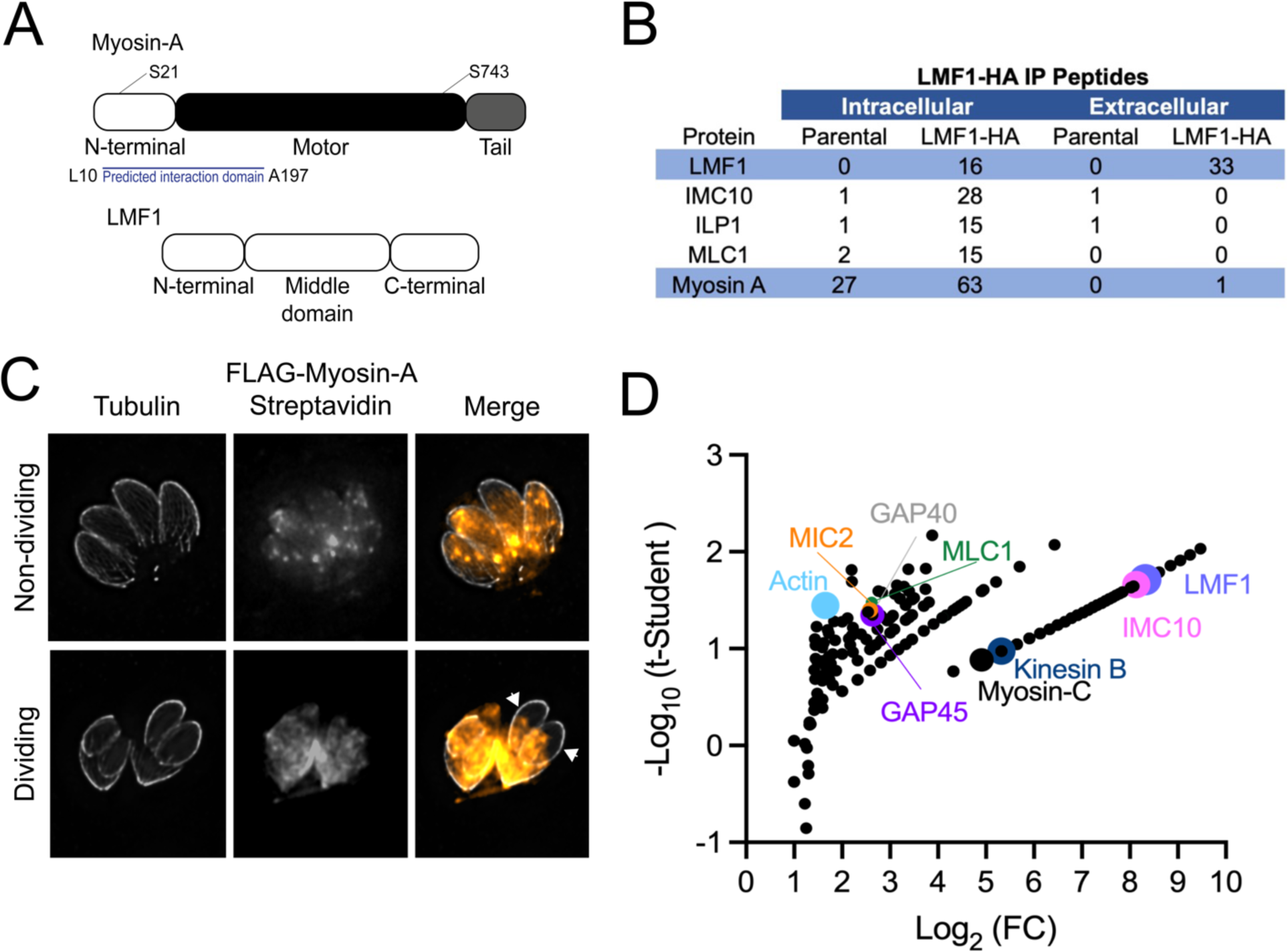
Myosin-A interacts with proteins involved in mitochondrial dynamics. A. Schematic of the three main domains in Myosin-A and LMF1, not to scale. The relative position of the two phosphorylation sites (S21 and S743) discussed in this work are marked. The region in MyoA found in the yeast two hybrid to interact with LMF1 is highlighted. B. Listed are the proteins identified in the IP of LMF1 from extracellular and intracellular parasites that are known to localize in the pellicle of the parasite. Table includes gene annotation and the number of peptides identified by mass spectrometry in both the control parental cell line (Δku80) and LMF1-HA. C. Immunofluorescence assays (IFA) of non-dividing and dividing intracellular FLAG-Myosin-A parasites stained for Tubulin (gray) and streptavidin-594 (orange). Arrows point at biotinylated proteins in the emerging daughter cells. D. Mass spectrometry analysis of FLAG-Myosin-A biotinylated proteins isolated using streptavidin beads. Highlighted are proteins known to interact with Myosin-A and proteins involved in mitochondrial dynamics, such as LMF1 (purple) and IMC10 (pink).

To further confirm the interaction between MyoA and LMF1, we used an established FLAG-tagged MyoA cell line (FLAG-MyoA) to perform Biotinylation by Antibody Recognition (BAR), a proximity labeling technique that has been validated for *Toxoplasma* (8). In this approach, intracellular parasites are fixed, permeabilized, and incubated with a primary antibody and a secondary horseradish peroxidase (HRP) conjugated antibody. Subsequently, the parasites are incubated with biotin-phenol and hydrogen peroxide to induce the biotinylation of proteins proximal to the HRP complexed protein of interest. To confirm the efficiency of the labeling, we performed a small-scale assay and performed IFA to detect biotinylated proteins by using an Alexa-Fluor conjugated Streptavidin. Curiously, there is not much enrichment of biotinylated proteins in the parasite’s plasma membrane when the parasites are intracellular (Fig. 1C). We co-stained with anti-tubulin, and it can be observed that in non-dividing parasites, the Streptavidin signal is distributed throughout the parasites, with several foci. When the parasites are dividing, we see a different pattern of staining, with a stronger and more concentrated signal in the basal end of the dividing parasites (Fig. 1C, arrows).

To identify proteins proximal to MyoA, we performed large-scale BAR assays with the FLAG-MyoA expressing strain and the parental strain as a control. Biotinylated proteins were captured using Streptavidin beads and analyzed by mass spectrometry. The criteria established to filter for the interactors are detailed in the Methods section. We generated a list of 144 putative proximal proteins (Fig. 1D) (Supplementary dataset S1). Among the proteins identified, we observed the presence of glideosome components such as GAP40, GAP45, and the myosin light-chain MLC1, as well as actin. We also observe the presence of other pellicle components, such as IMC proteins (IMC14) and the microtubule-associated proteins SPM1 and SPM2 (23), a calcium-dependent protein kinase CDPK2a and a Kinesin B, known to be located at the subpellicular microtubules (24). Surprisingly, we could identify peptides corresponding to both LMF1 and IMC10, showing that using this technique, we could capture the complex that tethers the mitochondrion to the pellicle. Thus, the mapping of MyoA protein-protein interactions suggests an association with proteins known to be involved in mitochondrial dynamics.

### Depletion of MyoA causes defects in mitochondrion morphology

To investigate the role of the MyoA motor in mitochondrial morphology, we took advantage of a previously established MyoA full knockout (25). This mutant strain has no defects in intracellular division but exhibits defects in egress and invasion (25, 26). As a control, we used a MyoA KO complemented with a wildtype version of MyoA (Complemented), which restores the egress phenotype (26). We stained parasites for F1B-ATPase to monitor the mitochondrion and acetylated tubulin to differentiate between dividing and non-dividing parasites and monitored the mitochondrial morphology in non-dividing parasites (Fig. 2). Based on the previous description of the mitochondrion; we classified the morphology as either lasso, sperm-like or collapsed (3, 5, 27, 28) (Fig. 2A). As expected, mitochondrion morphology in the complemented parasite strain was mostly in the lasso conformation (90±2.82%), which is like what we and others have reported for wildtype strains (3, 5). In contrast, MyoA KO parasites showed drastic changes in mitochondrion morphology, with most of the cells presenting sperm-like shaped mitochondrion (84.0±5.7%) and the appearance of collapsed mitochondrion (5.0±1.4% vs 0 in the complemented). To confirm that this phenotype was specific to the mitochondrion, we monitored the morphology of other organelles, including the apicoplast, rhoptries, endoplasmic reticulum, and IMC. No morphological defects were observed in any of these organelles in either knockout or complemented parasites (Supplemental Figure S2).

**Figure 2.**
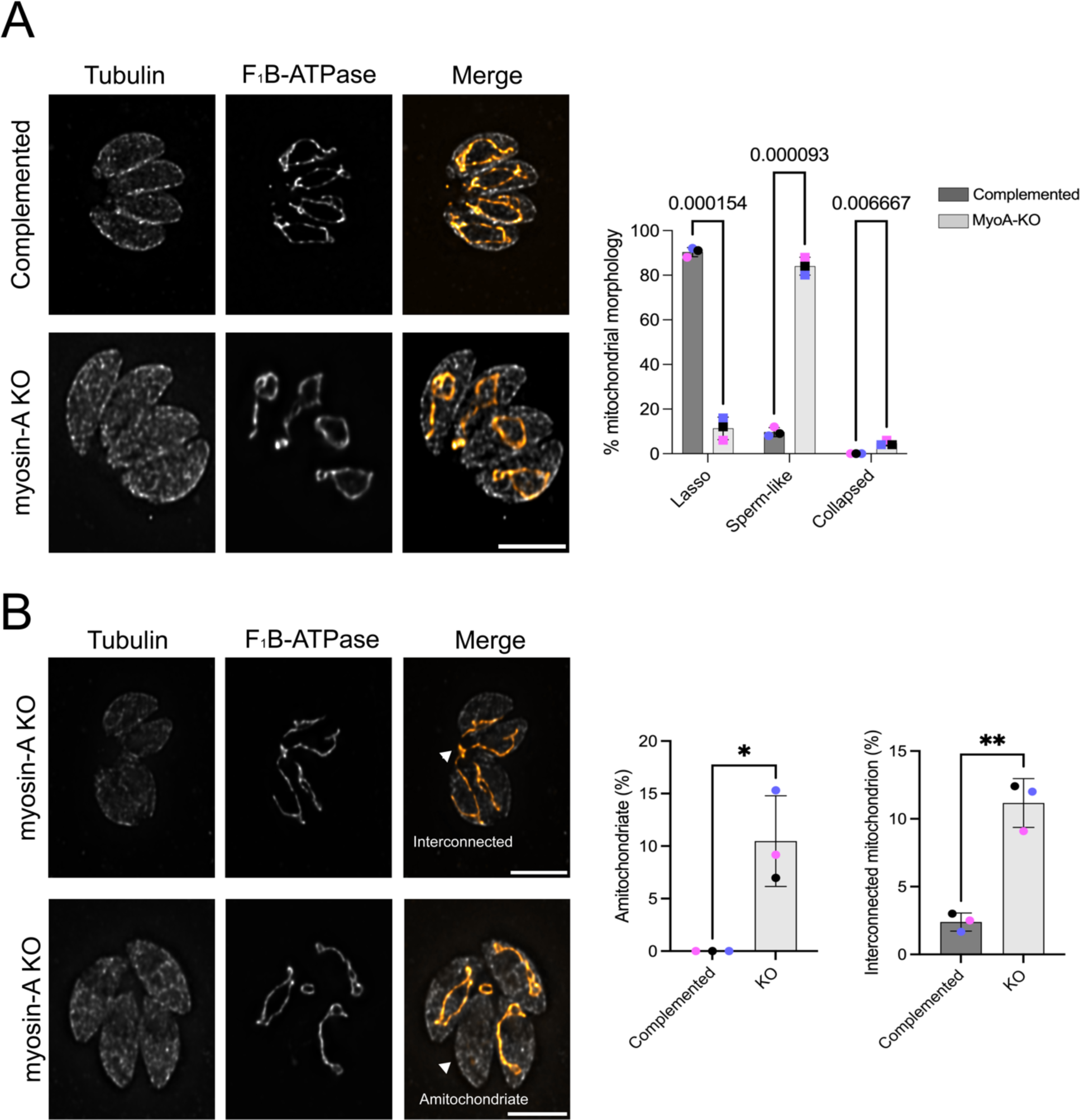
Depletion of Myosin-A affects mitochondrion morphology in intracellular parasites. A. Intracellular parasites of the FLAG-Myosin-A and myosin-A KO were grown in fibroblasts for 16 hours, then stained for Tubulin (gray) and F_1_B-ATPase, a mitochondrial marker (orange). Scale bar: 5 μm. Bar graphs represent percentage of parasites with each of the three morphologies in the presence or absence of Myosin-A. Data are means ± s.d. of three replicates; at least 150 non-dividing parasites were counted. P values are from two-way ANOVA with multiple unpaired *t*-tests. P <0.0001 significant; P>0.999 not significant. B. IFA of Myosin-A KO parasites showing parasites without mitochondrion and parasites with interconnected mitochondrion (arrows). Bar graph represents the percentage of parasites with each of these two phenotypes. ** P<0.01; * P<0.05; * P<0.05 (unpaired t-test, Two-tailed).

Besides the loss of the lasso morphology, the MyoA knockout strain shows phenotypes that might be related to defects in partitioning the organelle during inheritance. In comparison to the complemented strain, we observe the appearance of parasites lacking mitochondrial material in the MyoA KO strain (9.2±4.3% vs. 0 in the controls) (Fig. 2B). Also remarkable is the presence of vacuoles in which the mitochondria of different parasites are interconnected (12±1.8% of parasites in the knockout vs 2.5±0.7% in the complements). To confirm that the mitochondrion-related phenotypes observed with MyoA are specific to this Myosin, we monitored mitochondrial morphology in the absence of Myosin G using a conditional knockdown strain and observed none of the aberrant morphologies seen in the MyoA knockout strain (Supplemental figure S3). We also did not observe mitochondrial morphology in a strain lacking the calcium-dependent protein kinase 3 (CDPK3), which regulates MyoA during egress and whose knockout shares egress and motility defects with the MyoA knockout (26) (Supplemental figure S3). Thus, MyoA appears to have specific roles in the morphology and inheritance of the mitochondrion independent of its roles in motility.

### A specific point mutation in the MyoA N-terminal affects mitochondrial morphology

The Y2H screen identified the N-terminal of MyoA (amino acids 10 to 197) as sufficient for the interaction with LMF1. The N-terminal region of MyoA has been determined to be important for protein stabilization and modulating motility (29). Three amino acids within this region (S20, S21, and S29) are known to be phosphorylated (30) and we previously identified S21 as a substrate for CDPK3 (26). CDPK3 phosphorylation of MyoA at S21 as well as at S743, which is within the motor domain, is important for MyoA function within parasite egress (Fig. 11A) (26). Using previously established and validated strains, we tested whether mutation of S21 within the LMF1 interaction domain (S21A) or S743 within the motor domain (S743A) affected the function of MyoA in mitochondrial morphology (Fig. 3). Surprisingly, the parasites expressing the MyoA S21A mutant exhibited mitochondrial morphological defects, with few parasites presenting the normal lasso mitochondrion (14.0±2.8%), and most of the parasites showing sperm-like shaped mitochondrion (76.0±2.8%) (Fig. 3A). By contrast, the MyoA S743A mutant parasite presented normal mitochondrial morphology with 86.5±12.2%) presenting lasso-shaped mitochondrion (Fig. 3A).

**Figure 3.**
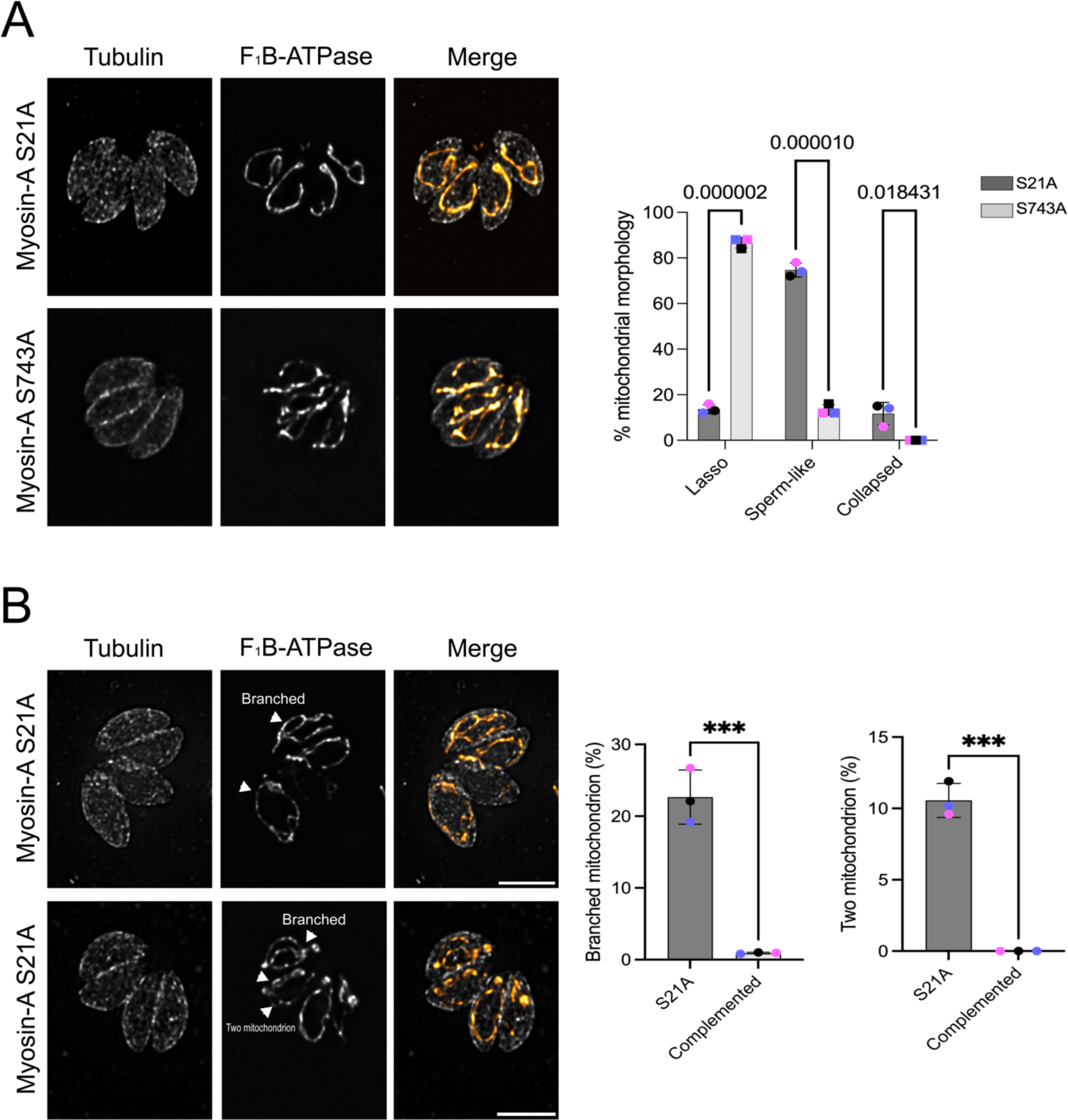
N-terminal phosphorylation of Myosin-A is important for mitochondrial morphology. A. Intracellular parasites expressing FLAG-Myosin-A S21A and FLAG-Myosin-A S743A were grown in fibroblasts for 16 hours and stained for tubulin (gray) and F_1_B-ATPase, a mitochondrial marker (orange). Scale bar: 5 μm. Bar graph represent percentage of parasites with each of the three morphologies in each of the two mutant strains. Data are means ± s.d. of three replicates; at least 150 non-dividing parasites were counted. P values are from two-way ANOVA with multiple unpaired *t*-tests. P <0.001, significant; P>0.999 not significant. B. IFA images of Myosin-A S21A parasites with arrows pointing at branched mitochondrion and at two independent mitochondria within one parasite. Data in bar graph are means ± s.d. of three replicates; at least 150 non-dividing parasites were counted. *** P<0.001.

A commonly observed secondary phenotype in the S21A mutant but not in the S743A one was single non-dividing parasites with an internally branched mitochondrion (Fig. 3B) that resembles dividing mitochondrion (22.1±3.8% in S743vs 1.0±0.1% in S743). Some individual MyoA S21A parasites also appear to have two independent mitochondria (10.2±0.9%) (Fig. 3B). Thus, our data suggest that residue S21, which is known to be phosphorylated and within the region of interaction with LMF1 (Fig. 1A), is important for MyoA’s role in mediating organellar morphology. To examine the specificity of this effect, we have also examined the positioning and morphology of other organelles, such as rhoptries, apicoplast, and endoplasmic reticulum in our mutant strains. Neither of the mutations affected the morphology of any of these organelles analyzed, corroborating the MyoA phenotypes observed are specific to the mitochondrion (Supplemental figure S4).

### Ultrastructure Expansion Microscopy (U-ExM) reveals Myosin A dynamic localization in dividing cells

MyoA is known to be located at the parasite’s pellicle (21, 31). Using Ultrastructure Expansion Microscopy (U-ExM), we revisited the localization of MyoA to increase the level of detail and observe the distribution of the protein within the cell with a higher resolution. Intracellular parasites expressing FLAG-tagged MyoA were expanded in a sodium acrylate hydrogel, and the cells were stained with anti-FLAG to visualize MyoA, anti-ATP beta subunit to monitor the mitochondrion, and with N-hydroxysuccinamide ester (NHS-Ester), which binds to all primary amines of proteins and can reveal cell structures with a high level of detail (32). In non-dividing parasites, we can observe that MyoA is distributed along the parasite’s pellicle/plasma membrane in a punctate pattern (Fig. 4A). No significant association with the mitochondrion is observed at this stage. In the late stages of division, when the daughter cells are fully formed, and the mitochondrion starts its entrance into the daughter cells, we detected a few spots corresponding to FLAG-MyoA surrounding the branching mitochondrion (Fig. 4A). These spots don’t overlay the anti-ATP ß-subunit staining, suggesting that these spots might be associated with the outer mitochondrial membrane (OMM).

**Figure 4.**
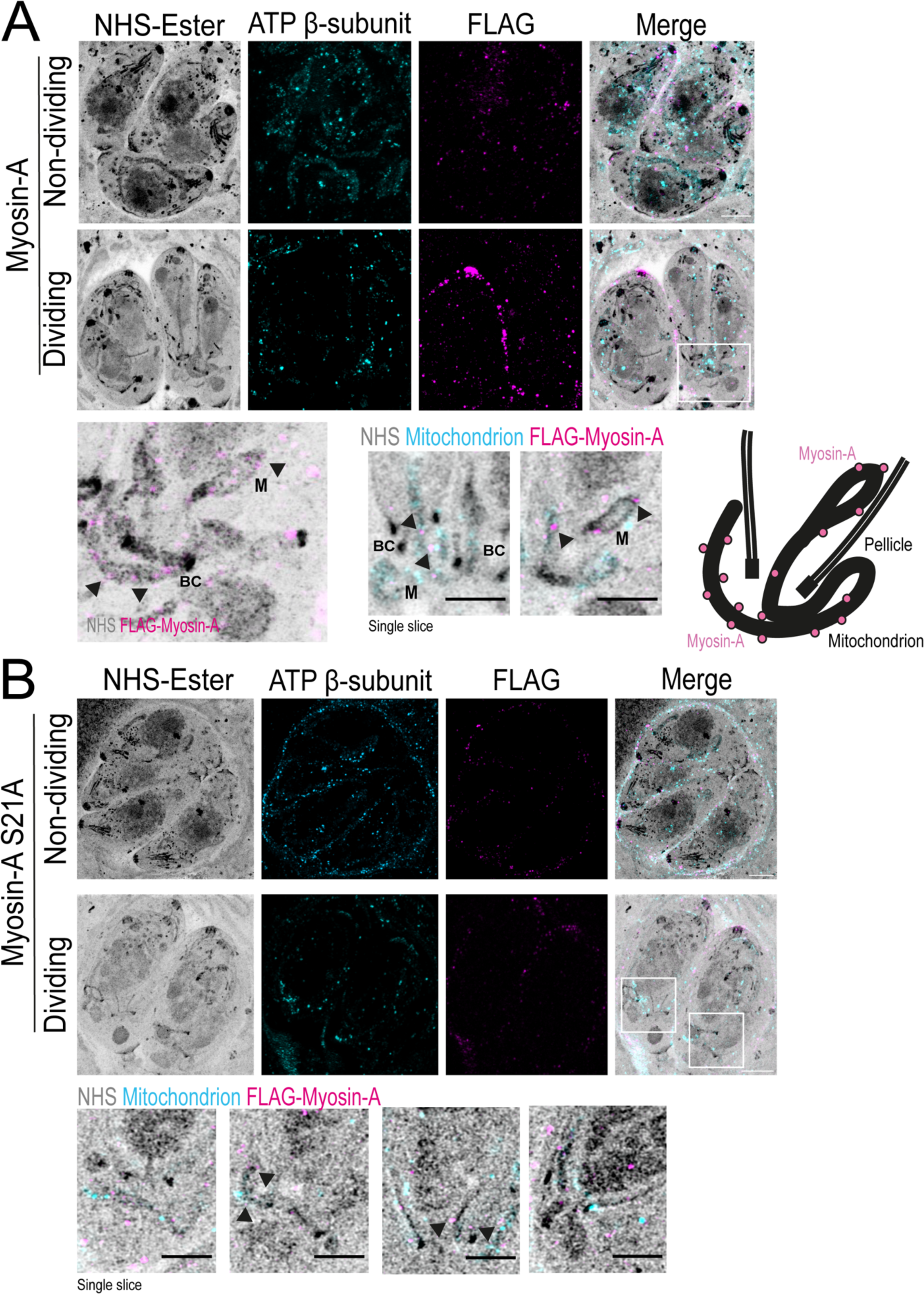
Expansion microscopy shows Myosin-A dynamic localization during cell division. A. U-ExM images of non-dividing and dividing intracellular parasites stained for total protein density (NHS-Ester), ATP β-subunit (mitochondrion, cyan), and FLAG-Myosin-A WT (magenta). The area in the white box in dividing parasites is magnified below to the left. To the right are single-slice images showing foci of Myosin-A near the parasite mitochondrion (arrows). BC = basal complex. Schematic to the right depicts the relative localization of the proteins and main structures, such as the pellicle, basal complex, mitochondrion, and Myosin-A (pink circles). B. U-ExM images of non-dividing and dividing intracellular parasites of the S21A mutant strain stained for total protein density (NHS-Ester), ATP β-subunit (mitochondrion, cyan), and FLAG-Myosin-A S21A (magenta). Bellow, single-slice images showing foci of Myosin-A near the parasite mitochondrion. M = mitochondrion. Scale bar in A and B: 5 μm.

As the S21A mutation affected mitochondrion morphology and it is within the LMF1-interacting domain (Fig. 1A), we decided to look if the protein would have the same pattern of mitochondrion localization during the late stages of cell division. The MyoA S21A mutants show the same localization as the wild type during interphase, with a punctate pattern spread all over the parasite’s periphery and extending to the parasite’s conoid (Fig. 4B, non-dividing). No clear mitochondrion localization was observed in this stage. However, during the late stages of division, we can observe some puncta corresponding to MyoA surrounding the branching mitochondrion, as observed for the wild type, suggesting that this mutation is not affecting its mitochondrial association (Fig. 4B).

### MyoA is involved in mitochondrial inheritance in *Toxoplasma*

The results described above confirm that MyoA plays a role in mitochondrial association with the periphery of the parasite. Moreover, we unexpectedly detected Myo in association with mitochondrial branches during parasite division, suggesting a possible role in mitochondrial inheritance. Accordingly, we explored the role of MyoA inheritance by monitoring mitochondria in dividing parasites in the MyoA KO and complemented parasites (Fig. 5). To narrow our analysis, we focused on parasites at a late stage of division when two daughter cells are completely formed. Furthermore, we only analyzed parasites with an elongated mitochondrion near the basal end to avoid biasing the results with collapsed mitochondrion, which is only seen in the KO parasites. Using these criteria, we quantitated the percentage of daughter parasites with an invading mitochondrial branch (Fig. 5A). The MyoA KO parasites showed a reduction of 60% of daughter cells presenting a mitochondrion branch (Fig. 5A). Similarly, the S21A mutants showed a reduction of 10% fewer daughter cells with a mitochondrion branch in comparison to the complemented strain. These results suggest that MyoA might participate in the process of directing the mitochondrion to the daughter cells and ensure proper mitochondrion inheritance in *Toxoplasma*.

**Figure 5.**
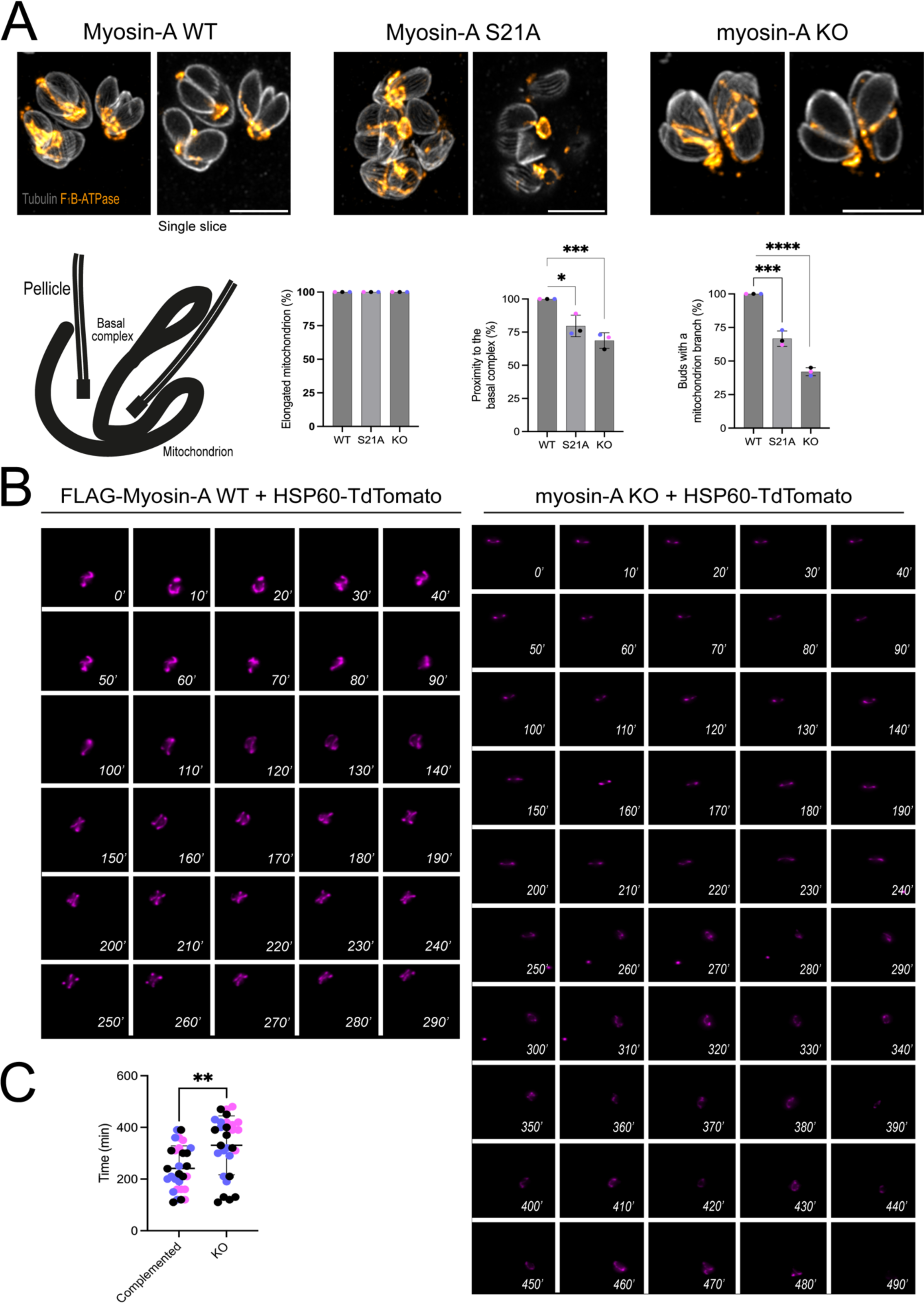
Myosin-A is critical for efficient mitochondrion inheritance. A. Representative IFAs of Complemented (Myosin-A WT), S21A, and Myosin-A KO intracellular parasites in the late stages of division. Parasites were stained for tubulin (gray) and F_1_B-ATPase (orange). For each strain, a representation of the maximum intensity projection of 35 Z-stacks is on the left and a single slice depicting the mitochondrion entering the daughter cell is on the right. Scale bar: 5 μm. Below, the scheme shows the normal phenotype expected for mitochondrion distribution. Graphs representing the percentage of elongated mitochondrion, proximity to the basal complex, and daughter cells with a mitochondrion branch. Data are means ± s.d. of 35 parasites per replicate, in three biological replicates. **** P<0.0001, *** P<0.001, * P<0.05 (unpaired t-test, Two-tailed). B. Time-lapse images of freshly invaded parasites during a cycle of division for the complemented or myosin-A KO parasites expressing HSP60-TdTomato. Images were acquired during 8 hours in intervals of 10 minutes per image acquired. The numbers in each image are the minutes from the beginning of imaging. C. Quantification of the average time for mitochondrion duplication in each condition. Data are mean ± SEM. A total of 30 parasites total per condition. **** P<0.0001, *** P<0.001, * P<0.05 (unpaired t-test, Two-tailed).

To better understand the role of MyoA in mitochondrion inheritance, we performed live-imaging cell microscopy. For this purpose, we transfected parasites (MyoA KO and complemented) with a construct that fuses the C-terminal portion of Hsp60 (including the mitochondrion-targeted signal peptide) to a TdTomato (2) and selected for positive clones with a fluorescent mitochondrion. Using these strains, we measured the time to duplicate and divide a mitochondrion after invasion. Imaging began 30 minutes post-infection, and single parasites were imaged every 10 minutes for 8 hours in total. In the complemented parasites, it is possible to observe that the mitochondrion is very dynamic in terms of internal movement (Movie m1). Parasites presenting a sperm-like mitochondrion at the start of imaging can reestablish the lasso shape in approximately 20-30 minutes of imaging (Movie m2). As these parasites start to prepare for the first round of division, the mitochondrion detaches from the pellicle (Fig. 5B, WT 30’), initiates elongation (60’), and divides into two newly formed daughter cells (Fig. 5B, WT 120’). Using the time-lapse videos, we calculated the average time to detect two individual mitochondria within each of the 30 parasites imaged per strain. For the complemented parasites, the average time to have two new mitochondria fully distributed among the daughter cells is 220±87.02 minutes. On the other hand, when we inspected the time for mitochondrion division in the MyoA KO parasites, we initially observed that this organelle is not as dynamic as what is observed in the complemented parasites (Movie m3). Upon infection, the mitochondrion stays in its initial shape for an average of 120 minutes. From this time point, the organelle starts its branching (Fig. 5B, KO 250’) and elongation process (Fig. 5B, KO 260’-270’), albeit with slower movements than what is seen in the control (Movie m4). The calculated average time to have two new mitochondria fully distributed among the daughter cells is 370±113.92 minutes. Interestingly, we commonly observed MyoA KO parasites in which the mitochondrion branches only start migrating into the daughter parasites after emerging from the mother (movie m#). Thus, the live cell imaging data, along with the U-ExM images, support the hypothesis that MyoA is implicated in mitochondrial inheritance in *Toxoplasma*.

### Conditional knockdown of LMF1 in the MyoA mutants exacerbates phenotypes

Previous studies have shown that LMF1 is not essential for parasite viability but affects parasite propagation in tissue culture (5). Similarly, the MyoA knockout parasites exhibit a slower rate of propagation (21, 25). As we have shown that MyoA and LMF1 interact and both affect the mitochondrial dynamics, we investigated the effect of disrupting both proteins simultaneously. For this purpose, we added a destabilization domain (DD) and a myc epitope tag at the C terminus of the endogenous LMF1 in the MyoA KO, complemented, and S21A strains. The DD targets the protein for degradation unless the stabilizing agent Shld1 is added to the growth medium (Fig. 6A). Once all strains were established, we validated localization and expression by IFA (Fig. 6A) and western blot (Fig. 6B). Immunofluorescence assays in the presence of Shld1 show that LMF1 is correctly localized in all strains (Fig. 6A). Interestingly, by IFA, we do not detect any LMF1 after 24 hours without Shld1 for any of the strains (Supplemental figure S5). To confirm that the DD domain, was not affecting the interaction between LMF1 and MyoA, parasites were grown in the presence of Shld1 for 48 hours and we performed co-immunoprecipitation using anti-myc beads (Fig. 6B). Using this technique, we confirmed that immunoprecipitation of LMF1 with myc beads brought down MyoA, both the WT type protein (complemented) and S21A mutant protein (Fig 6B).

**Figure 6.**
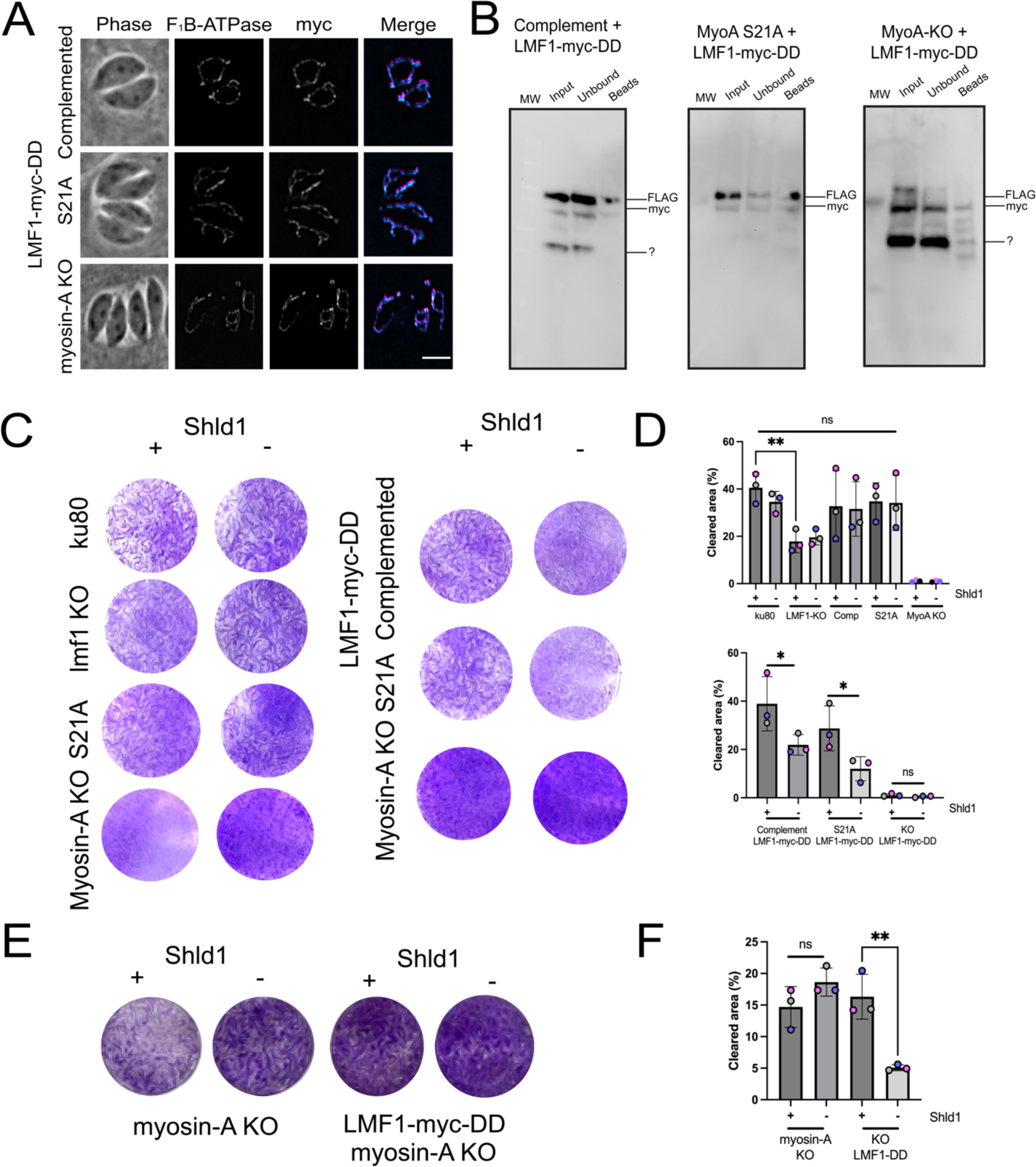
Establishment and growth of parasite strains with a conditional expression of LMF1. A) Representative IFA of parasites expressing LMF1-myc-DD showing localization of LMF1 in the presence of Shld1. Parasites were stained for F_1_B-ATPase (cyan) and myc (magenta). Scale bar: 5 μm. B) Western blot of co-immunoprecipitation (co-IP) of MyoA mutants (complement, S21A, and KO) expressing LMF1-myc-DD. LMF1 was immunoprecipitated with anti-Myc conjugated beads and probed with anti-Myc for LMF1 and anti-FLAG for MyoA. (?) is a non-specific band. D. Quantification of the total cleared area per well for either parental or knockdown cell lines grown in the presence (+) or absence (-) of Shld1 after a 5-day incubation period. E. Representative plaque assay of myosin-A KO cell line expressing LMF1-myc-DD in the presence (+) or absence (-) of Shld1 for 10 days. F) Quantification of the total cleared area per well for either parental or knockdown cell lines grown in the presence (+) or absence (-) of Shld1. Plaque assays were performed in three biological replicates. Data are means ± s.d.. Statistical analysis is a Two-tailed unpaired t-test, ** P<0.01; * P<0.05; ns not significant (P>0.05).

With the validated strains in hand, we evaluated parasite growth of the parental and the LMF1-myc-DD expressing strains in the absence or presence of Shld1 for five days using plaque assays (Fig. 6C). We used the parental Λ1ku80 and an lmf1 knockout strain as controls (5). We first assessed plaque formation in the parental strains (e.g. not expressing LMF1-myc-dd) with and without Shld1 (Fig. 6C). As expected, the absence of Shld1 does not affect the growth of any of these strains. Nonetheless, we confirmed that the LMF1 KO strain has a reduction in propagation based on the clearance of the host cell monolayer (41.6±6.3% for ku80 vs 16.3±4.7% for lmf1KO) (Fig. 6C). Both the MyoA complemented and S21A mutant showed similar plaquing efficiency as the ku80 strain (Fig. 6C). At five days plaques were not visible for the MyoA KO strain, so we allowed the plaques to develop for 10 days as to be able to quantitate propagation (Fig. 6E).

We then assessed the growth with and without Shld1 for the LMF1-myc-DD strains (Fig. 6D). LMF1 knockdown in the MyoA wildtype reduced the level of clearing of the host cell monolayer (19.9±4.3% without Shld1 vs 31.0±14.5% with) (Fig. 6D). In a similar fashion, LMF1 knockdown reduced plaque clearing in the S21A background (14.5±4.9% without Shld1 vs 26.02±9.32% with Shld1) (Fig. 6D). The reduction in propagation for these two strains upon loss of LMF1 is consistent with the effect of loss of LMF1 in the wildtype background (Fig. 6C). In the MyoA KO background after 10 days of growth, we observed a pronounced reduction in propagation upon knockdown of LMF1. In the presence of Shld1, the MyoAKO/LMF1mycDD strain cleared 14.3±3.6% of the host cell monolayer. The same strain disrupted 4.9±0.5% of the monolayer when grown without Shld1, which resulted in the loss of LMF1 (Fig. 6E). Thus, the knockdown of LMF1 in a complemented background reduces propagation by 34% (Fig. 6D, +Shld1 vs – Shld1) and by 65% in a MyoA KO background (Fig. 6E, +Shld1 vs – Shld1). Together, these results indicate that the depletion of LMF1 in the MyoA KO background affects parasite replication and that loss of both LMF1 and MyoA exacerbates the propagation phenotype seen in each of the single knockout strains.

### Conditional knockdown of LMF1 in the MyoA mutants affects parasite mitochondrial distribution and morphology

As both LMF1 and MyoA are involved in mitochondrial dynamics and mitochondrial inheritance during cell division, we evaluated the combined effects of simultaneously disrupting MyoA and LMF1 expression. We infected coverslips with all LMF1-myc-DD strains in the presence or absence of Shld1 for 24 hours (Fig. 7). After this period, we examined mitochondrion morphology by IFA, after staining parasites with anti-IMC10 and anti-ATOM40. In the wildtype MyoA background, resulting in a loss of LMF1 decrease in lasso shape (5.7±3.45% -Shld1 vs. 77.5±2.64% +Shld1) and an increase in sperm-like (61.3±4.452% -Shld1 vs. 21±2.02% +Shld1) and collapsed-shaped mitochondrion (29±3.78% -Shld1 vs 1±0.76% +Shld1) (Fig. 7A). Again, these results are very similar to what was observed in the lmf1 knockout strain (Jacobs et al, 2020). When we inspected the mitochondrion morphology in the S21A/LMF1mycDD strain, it was possible to observe some parasites with a lasso mitochondrion (10.1+3.4%) in the +Shld1 condition, but the proportion of parasites with sperm-like (79.6±6.3%) and collapsed (5.2±2.0%) mitochondrion was very similar to what we’ve observed previously for this mutant (Fig. 7B). Interestingly, in the absence of Shld1 and thus LMF1, we can still detect some lasso (14.6+4.9%), but notably there is a marked decrease in parasites with sperm-like (52.0±3.0% -Shld1 vs 79.6±6.25% +Shld1) and an increase in the collapsed mitochondrion (33.4±7.85% - Shld1 vs 5.2±7.7 +Shld1), showing that the absence of LMF1 in the MyoA S21A exacerbates organelle collapse (Fig. 7B). In the MyoA KO/LMF1mycDD strain grown with Shld1, we see the same severe mitochondrial morphology defects as in the original MyoA KO (Figs. 2 and 7). By contrast, when MyoA KO/LMF1mycDD is grown without Shld1, we observe a significant decrease in sperm-like (46.4±4.2% -Shld1 vs 72.9±10.61% +Shld1) and an increase in the collapsed mitochondrion (38.3±9.05% vs 25.6±9.67 in the +Shld1) (Fig. 7C). The number of cells showing a lasso shape does not change in either condition. Altogether, we confirm that decreasing the levels of LMF1 in the MyoA mutant backgrounds affects mitochondrial morphology, resulting in increased collapsed mitochondrion.

**Figure 7.**
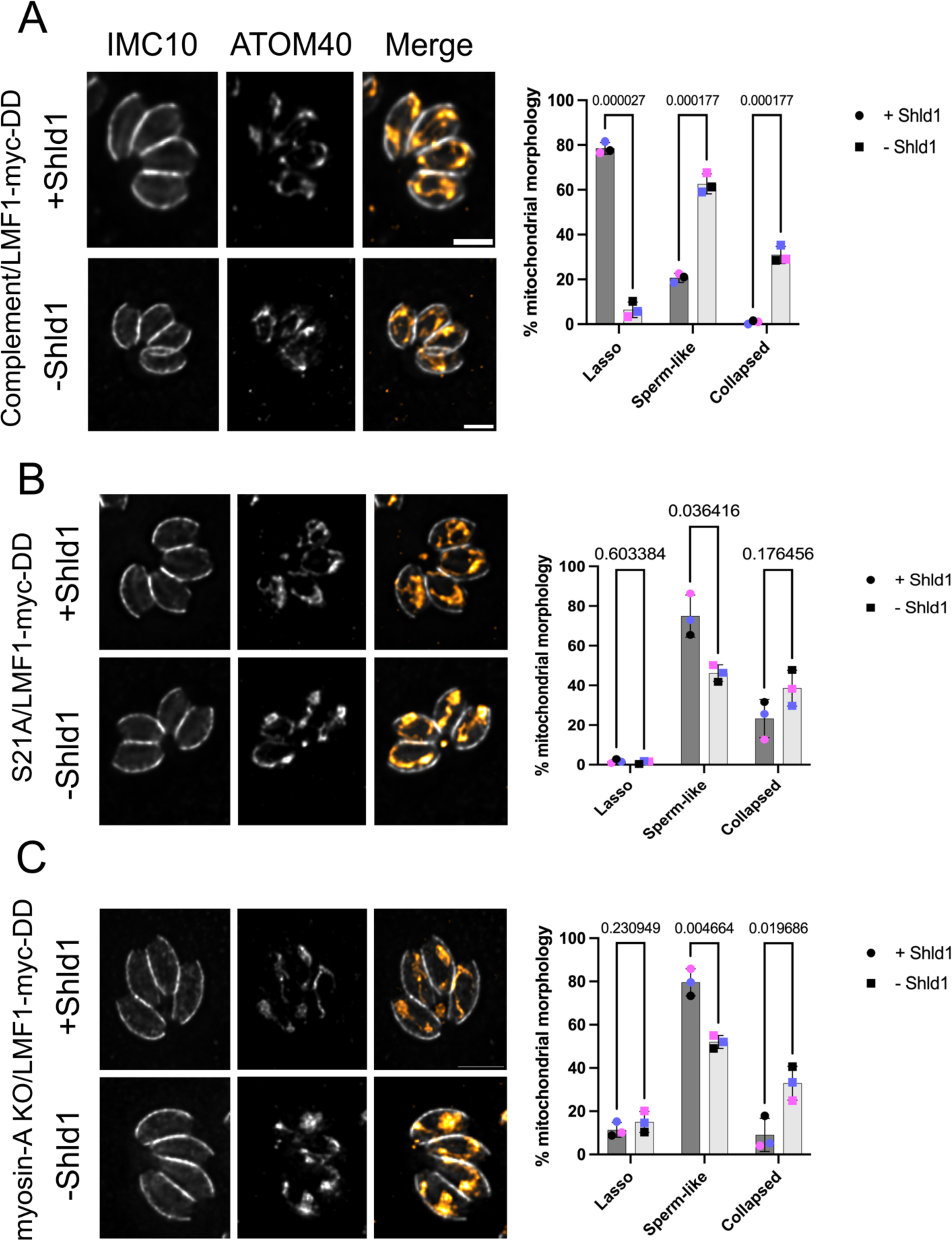
LMF1 conditional knockdown affects mitochondrion morphology in Myosin-A mutant intracellular parasites. Shown are IFA images of intracellular Complement (A), Myosin-A S21A (B), and myosin-A KO (C) parasites expressing LMF1-myc-DD. Parasites were grown in the presence (+) or absence (-) of Shld1 for 24 hours and stained for IMC10 (gray) and ATOM40, a mitochondrial marker (orange). Scale bar: 5 μm. Bar graphs represent the percentage of parasites with each of the three morphologies for each condition. Data are means ± s.d. of three replicates; at least 150 non-dividing parasites were counted. *q* values represented in each graph. P values are based on two-way ANOVA with multiple unpaired *t*-tests. P <0.0001 significant; P>0.999 not significant.

One of the defects observed in the MyoA KO cell lines is the presence of cells lacking mitochondrial material. We would expect that by controlling LMF1 protein levels, we would increase the number of vacuoles with abnormal mitochondrial distribution. When in the absence of Shld1, it is indeed possible to observe vacuoles containing abnormal mitochondrion distribution. We inspected all the conditions, looking for vacuoles where i. we could observe mitochondrial material concentrated outside of the parasites and amitochondriate parasites, ii. parasites with the mitochondrial material stuck inside of a single parasite, while the other cell within the same vacuole completely lacks mitochondrion. We can find vacuoles where there is a single mitochondrion in one parasite, whereas the other parasites lack mitochondrion material (Fig. 8A, arrows). To sum up this phenotype, we decided to name it abnormal mitochondrial distribution. When in comparison to the parasites kept in the presence of Shld1, there are no significant differences between the conditions, both for 24 hours (4.8±1.35% vs. 7.6±1.43% in the -Shld1) and 48 hours (8.9±0.78% vs. 10.5±1.26% in the -Shld1) (Fig. 8A).

**Figure 8.**
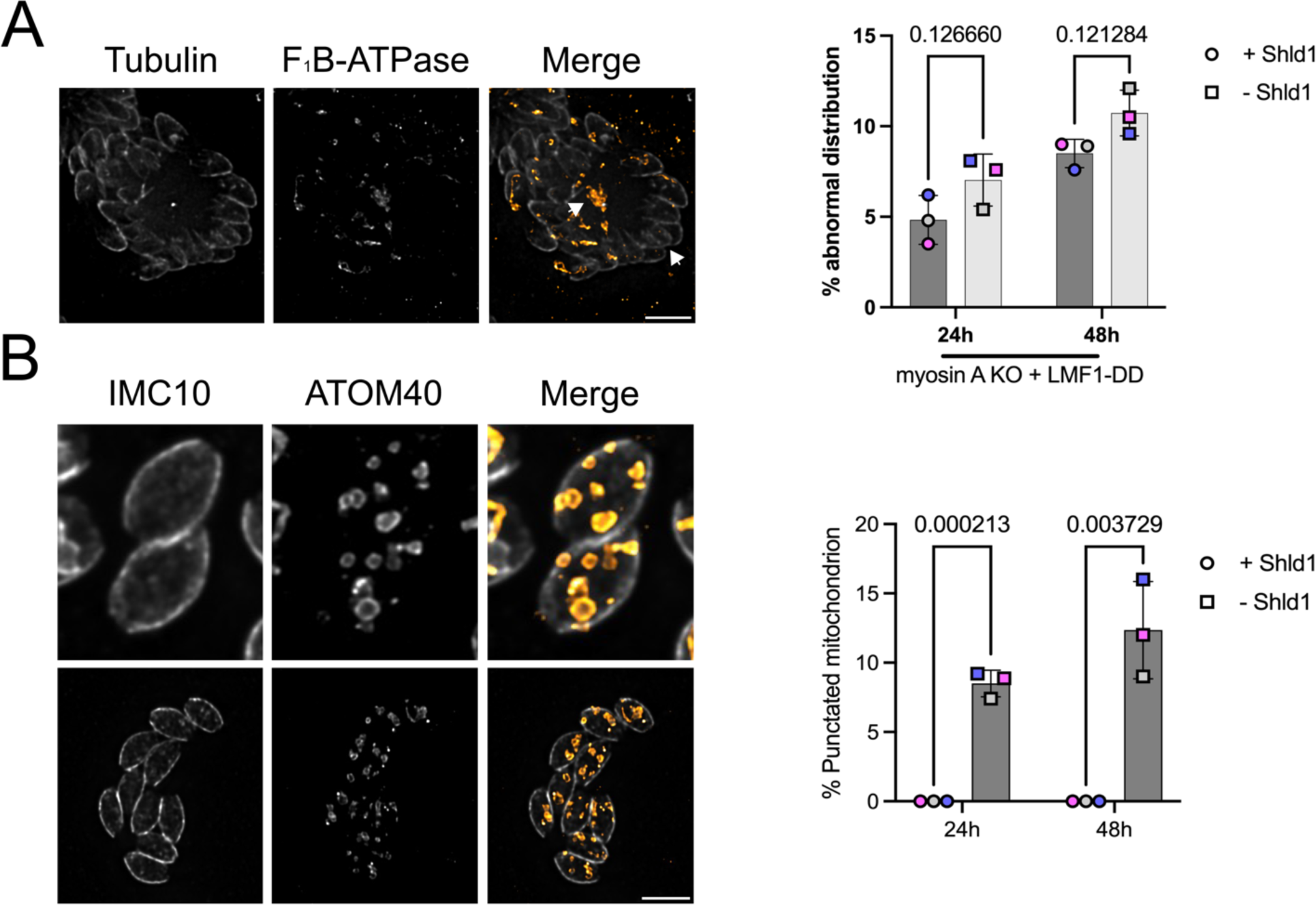
LMF1 knockdown in Myosin-A knockout background results in mitochondrial fragmentation. A. IFA of myosin-A KO parasites carrying LMF1-myc-DD grown in the absence of Shld1 stained for Tubulin (gray) and F_1_B-ATPase (orange) showing the accumulation of mitochondrion in the residual body (arrows). Bar graph represents the percentage of parasites with abnormal distribution of mitochondria with and without Shild1. Data are means ± s.d. of three replicates with at least 100 vacuoles counted per condition. P values are from two-way ANOVA with multiple unpaired *t*-tests, P <0.0001 significant; P>0.999 not significant. B. IFA of myosin-A KO parasites carrying LMF1-myc-DD in the absence of Shld1 stained for IMC10 (gray) and ATOM40 (orange) showing fragmented mitochondrion through the parasites. Scale bar in A and B: 5 μm. Bar graph represents the percentage of parasites with fragmented mitochondria. Statistics same as A.

Strikingly, the simultaneous disruption of both MyoA and LMF1 results in a remarkable phenotype not observed in either the single MyoA or LMF1 mutant strains. In 8.9±1.0% of MyoA KO/LMF1-myc-DD parasites grown without Shld1 for 24 hours, we detected multiple pieces of mitochondria within a single parasite (Fig. 8B). This phenotype becomes more common after 48 hours without Shld1 (12.0±3.5%) (Fig. 8B). As we stained the parasites with anti-ATOM40, which detects the outer mitochondrial membrane, we can infer that these are individual pieces of organelle containing a closed outer mitochondrion membrane. Remarkably, this phenotype is not present in any of the single mutant strains we inspected or in the same strain grown with Shld1, indicating that it is unique to the simultaneous lack of both LMF1 and MyoA.

### Disrupting actin, but not microtubules, affects mitochondrion morphology

Given the role of MyoA in mitochondrial dynamics, we sought to determine if actin is also involved in this process. To test this possibility, we monitored mitochondrion morphology in parasites treated with the actin destabilizer Cytochalasin D. Specifically, we allowed parasites to grow for 12 hours before adding either 1μM Cytochalasin D or DMSO for 16 hours (Fig. 9). IFAs using antibodies against tubulin and mitochondrion showed that actin depolymerization substantially affected the parasite’s mitochondrion morphology (Fig. 9A). In vacuoles with two or four parasites, nearly all exhibited sperm-like mitochondrion restricted to the parasite’s periphery and aligned with the subpellicular microtubules (Fig. 9A). Interestingly, in vacuoles with 16 or more parasites, it’s possible to observe parasites lacking mitochondrion mitochondrial material (Supplemental figure S6).

**Figure 9.**
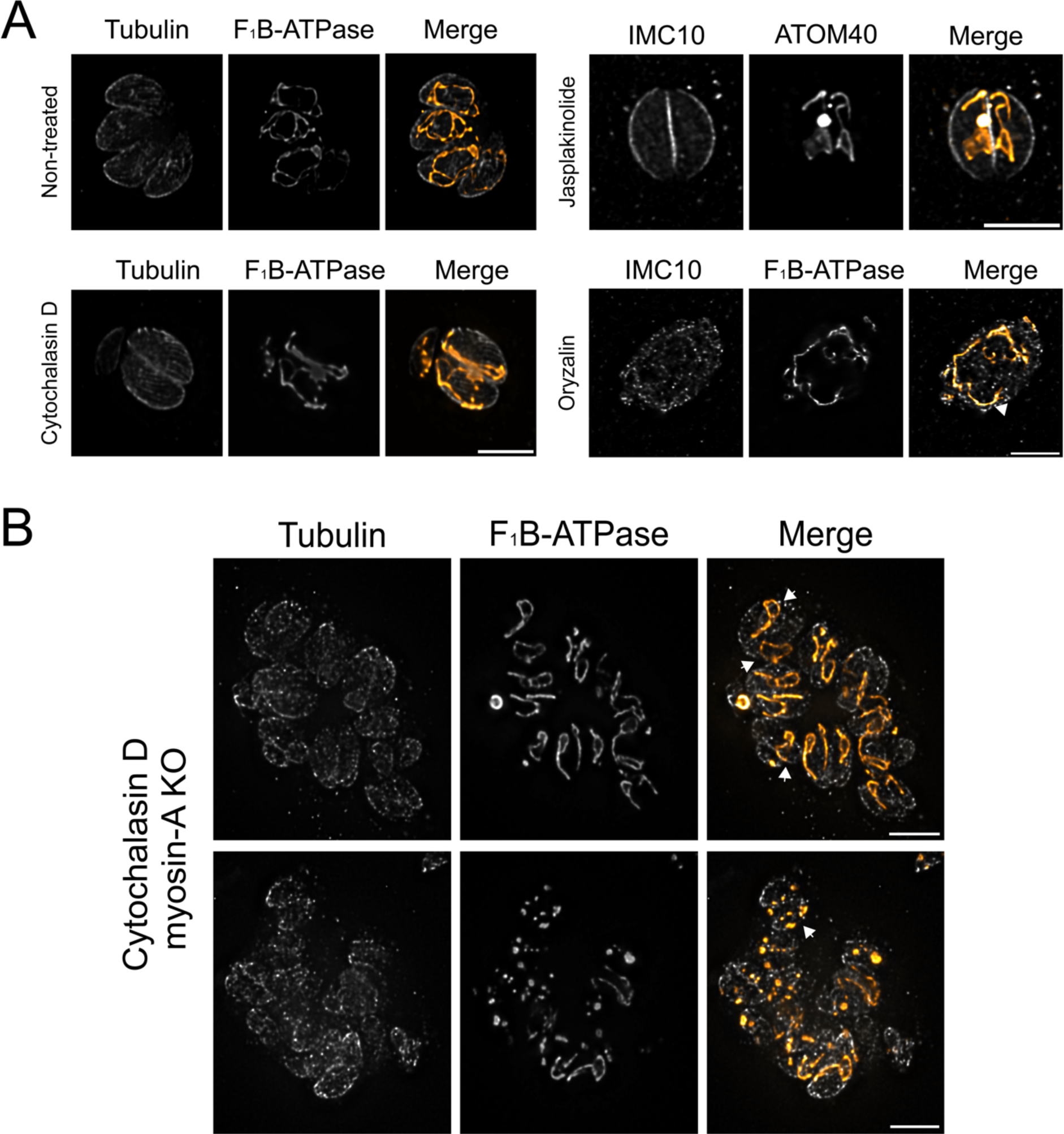
Actin is critical for mitochondrion morphology. A. IFA of parasites grown in the presence of 1 μM of Cytochalasin D, 1 μM Jasplakinolide, 1 μM Oryzalin, or DMSO (non-treated). Parasites were stained for Tubulin (gray) and F_1_B-ATPase (orange). B. IFA of myosin-A KO parasites growing the presence of Cytochalasin D (1 μM) and stained for Tubulin (gray) and F_1_B-ATPase (orange). Arrows point at mitochondria that appear outside of the parasites (top right) and fragmented mitochondrion (bottom right).

Given the fact that destabilizing actin affected mitochondrion morphology, we sought to analyze if actin stabilization would also affect organelle morphology. For this purpose, we treated intracellular parasites with 1μM Jasplakinolide for 2 hours. In contrast to the cytochalasin D treatment, we did not observe parasites without mitochondrial material in larger vacuoles in the Jasplakinolide-treated samples (Supplemental figure S6). Nonetheless, similarly to Cytochalasin D, Jasplakinolide affected the parasite’s mitochondrion morphology (Fig. 9B). Interestingly, instead of a sperm-like shaped mitochondrion, the organelle appears to be more branched and accumulated in the middle of the cell. As a control, we have analyzed mitochondrion morphology upon microtubule disruption with Oryzalin. As previously described (2, 3), we observed that Oryzalin severely disrupts the parasite’s morphology. However, it’s still possible to observe contact between the mitochondrion and the parasite periphery, showing that microtubules are not important either for tethering the mitochondrion to the pellicle or for mitochondrion positioning (Fig. 9C).

Our previous results showed that the lack of both MyoA and LMF1 results in severe mitochondrion morphological and inheritance defects, including mitochondrion fragmentation. As actin dynamics exert an important role in mitochondrial dynamics, we decided to look at the effects of actin depolymerization combined with the loss of MyoA. Accordingly, we treated MyoA KO parasites with 1μM of Cytochalasin D and analyzed mitochondrial morphology by IFA (Fig. 9D). For the most part, addition of Cytochalasin D to the MyoA KO parasites did not affect the mitochondrion beyond what is already seen with this strain (Fig. 2 and Fig. 6D). Interestingly, we observe mitochondria that appear to be outside the parasite’s cytoskeleton (Fig. 9B, top right with arrows). This positioning could be the result of the organelle remaining outside of the parasites, probably during division. Another interesting characteristic observed in the Cytochalasin D treated MyoA KO parasites was fragmented mitochondrion distributed in spots inside of the cell (Fig. 9B bottom right with arrows). This phenotype resembles what we detected for the MyoA KO parasites upon LMF1 knockdown. Altogether, our results show that the actin network is important for proper mitochondrion morphology.

### Toxoplasma mitochondrion interacts with actin filaments in vitro

Yeast mitochondria are known to interact with actin filaments in vitro (13). The interaction between yeast mitochondria and actin is ATP-sensitive, reversible, and dependent on mitochondria-associated proteins (13-15, 17, 33). Given our observation that both actin and a myosin motor are needed for mitochondrial dynamics, we set out to investigate whether *Toxoplasma* mitochondrion could interact with actin in vitro. To achieve this goal, we adapted an established method to enrich mitochondrial fraction with an intact outer membrane using digitonin as an agent, which is known to selectively fractionate membranes in a concentration-dependent manner (34) (Fig. 10A). To facilitate monitoring mitochondria in the recovered fractions we used a strain expressing the enzyme superoxide peroxidase 2 tagged with an N-terminal GFP (SOD2-GFP) (35). Extracellular parasites were treated with either 0.1% or 0.05% of digitonin for 5 minutes, and we then separated the pellet and supernatant by centrifugation, and the resulting fractions were analyzed by immunoblotting (Fig. 10B). Blots were probed with anti-ATOM40 (outer mitochondrion membrane) and anti-GFP (to detect SOD2, inner mitochondrial membrane). As controls, we probed for eIF2 as a cytosolic marker, ACP to detect apicoplast contamination, and IMC and Tubulin to detect cytoskeleton contamination. In both digitonin treatments, we could observe that the cytosolic marker eIF2 is present only in the supernatant (SN), showing a successful separation of cytosolic and organellar material (Fig. 10B). Mitochondria appear to be enriched with the membrane fractions as most of the GFP signal is within the pellet. Importantly, the OMM appears to be intact, with only a small portion of the ATOM40 signal presented in the supernatant. Based on the immunoblotting, we can still detect contamination with the parasite’s cytoskeleton (IMC and Tubulin signal presented in the pellet fraction). It is also possible to detect the apicoplast in these fractions based on the ACS signal. We then performed the fractionation assays using the LMF1-HA, FLAG-MyoA, and FLAG-MyoA(S21A) strains to test whether LMF1 and MyoA would be detected in the mitochondrion-enriched pellet. In the western blots probed for both anti-HA and anti-FLAG, we can observe that the majority of both LMF1 and MyoA are detected in the pellet fraction, together with the mitochondrion marker ATOM40 (Fig. 10C).

**Figure 10.**
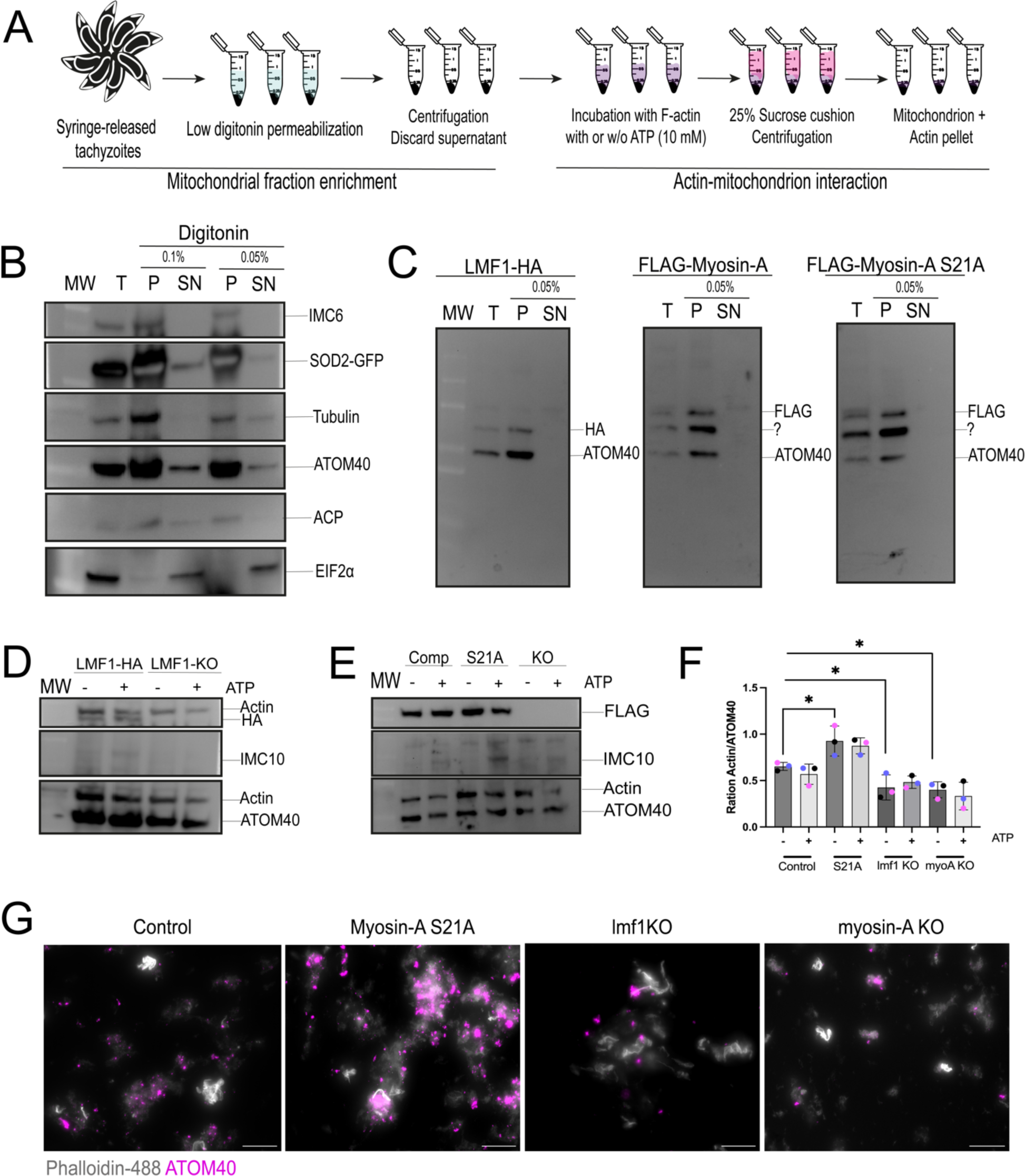
Actin and mitochondrion interact in vitro. A. Schematics of the procedure to extract mitochondrion-enriched fractions from intracellular parasites and perform the actin-mitochondrion interaction assy. B. Western blots of SOD2-GFP parasites subjected to organelle enrichment protocol. Parasites were treated with 0.1% or 0.05% digitonin. T: total extracts; P: pellet after digitonin fraction; SN: supernatant. Western blot membranes were probed for the presence of IMC (anti-IMC6), microtubules (anti-Tubulin), mitochondrion OMM (anti-ATOM40), apicoplast (anti-ACP), and cytosol (anti-EIF2α). SOD2-GFP (mitochondrion soluble marker) was tracked using anti-GFP antibodies. C. Western blots of fractions harvested from parasites expressing LMF1-HA, FLAG-Myosin-A and FLAG-Myosin-A S21A treated with 0.05% digitonin. The membranes were probed using antibodies against HA (LMF1) and FLAG (Myosin-A). Antibodies against ATOM40 were used to probe for mitochondria. The question mark points at a non-specific band. D and E. Mitochondria-enriched vesicles from LMF1-HA and lmf1-KO parasites (D) or myosin-A WT, S21A, and KO parasites (E) were incubated with F-actin in the presence or absence of ATP (10mM). Membranes were probed for the presence of LMF1 (anti-HA in D) or Myosin-A (anti-FLAG in E). Membranes were also probed for the presence of actin (anti-actin), IMC (anti-IMC10), and mitochondrion (anti-ATOM40). F. Quantification of actin/ATOM40 ration in the presence (+) or absence (-) of ATP. Quantification was performed by densitometry using FIJI. Data are means ± s.d. of three replicates. * P<0.05, unpaired t-test, Two-tailed. Each colored dot represents a different replicate. G. Mitochondrion-enriched fractions isolated from all the cell lines were stained with anti-ATOM40 (magenta), incubated with Phalloidin-488 labeled actin filaments (gray), and observed by fluorescence microscopy.

After confirming the enrichment of a mitochondrial fraction, we performed the mitochondrion-actin interaction assays (Fig. 10A). For this purpose, we incubated the enriched fractions with actin filaments in the presence and absence of ATP for 30 minutes, sedimented the organelles with a sucrose cushion followed by centrifugation. The sedimented fractions were analyzed by western blot probing for ATOM40 and actin antibodies. To confirm that these fractions carried mostly mitochondria and that the sedimented actin wasn’t bound to the pellicle, we also probed the blots with an antibody against IMC10. Our co-sedimentation assays showed that enriched mitochondrion fractions can interact with actin filaments (Fig. 10D and E). Another observation is the fact that after the sucrose cushion, apparently no IMC10 can be detected, showing that this protocol can help in increasing the mitochondrion enrichment. Importantly, we do not detect any tubulin or ACS (apicoplast) in the sedimented fraction, indicating that we have minimal if any, cytoskeleton and apicoplast contamination (Supplemental figure S7). Nonetheless, the presence of LMF1-HA (Fig. 10D) and FLAG-MyoA (Fig. 10E) can still be detected even after the sucrose cushion, showing that these proteins continue to enrich with the mitochondrion after the sucrose sedimentation. We quantitated results from triplicate experiments and expressed the data as the ratio of actin to ATOM40 signal (Fig. 10F). Interestingly, ATP is not able to inhibit binding, which shows that actin-mitochondrion co-sedimentation is ATP-insensitive (Fig. 10F). We performed the mitochondrion enrichment protocol in the LMF1-KO, MyoA-KO, and S21A and used these fractions to evaluate the actin co-sedimentation in these conditions. Compared to our controls, we can observe a reduction of 30% in co-sedimentation when parasites lack LMF1 (ratio 0.4±0.1) (Fig. 10F). Similarly, depletion of MyoA also affected actin binding to the mitochondrion (0.4±0.09), comprising ∼33% of the actin bound in the WT mitochondrion (Fig. 10F). Thus, co-sedimentation of LMF1 and MyoA is dependent on both LMF1 and MyoA. Interestingly, parasites carrying the mutated MyoA S21A showed an increase in actin sedimentation approximately 40% more (0.9±0.2) in comparison to the WT.

To corroborate these findings and visualize a plausible mitochondrion-actin interaction, we imaged the sedimented fraction using immunofluorescence. Briefly, we pre-stained the mitochondrion-enriched fraction using the anti-ATOM40 antibody and the actin filaments using Phalloidin-488, which were then incubated together and ran through the sucrose cushion. The experiments were done without ATP, as it did not affect binding. Samples were immobilized in mounting media and observed by fluorescence microscopy (Fig. 10G). In all the conditions, we can visualize mitochondria bound to pre-formed actin filaments. We can conclude from this series of experiments that *Toxoplasma* mitochondrion physically interacts with actin, and this interaction is partially dependent on MyoA and LMF1.

## DISCUSSION

Organelles are highly dynamic entities with the ability to change their morphology and positioning in a fast and efficient way (36). Organellar movement is of particular importance during cell division. In particular, the movement of mitochondria within the cell is an important step in cell division because the efficient partitioning of mitochondria guarantees proper mtDNA segregation and mitochondrial function within daughter cells (37). Mitochondrial segregation is particularly important in cells carrying a single mitochondrion. In this study, we focused on the proteins that collaborate in mitochondrion distribution in the parasite *Toxoplasma*. Importantly, we confirmed a putative interaction between the tethering protein LMF1 and MyoA. Although the role of MyoA in parasite motility has been well characterized (21), the role of this unconventional motor in organelle segregation has not been previously reported.

*Toxoplasma* possesses a repertoire of 11 myosins (38). MyoA belongs to the unusual Class XIVa myosins and is the most studied Myosin in apicomplexans because of its role in driving gliding motility (21) and its potential as a drug target against toxoplasmosis (39, 40). In *Toxoplasma*, this motor consists of a MyoA heavy chain bound to two light chains, MLC1 (41) and either ELC1 or ELC2 (42). Together with the glideosome-associated proteins GAP40, GAP45, and GAP50, MyoA forms part of a complex named glideosome (43, 44). This complex is immobilized on the IMC surface (42), generating a driving force that promotes parasite motility (45). In the proposed model for gliding motility, actin filaments present in between the IMC and the plasma membrane are connected to ligands through connector proteins called GACs (glideosome-associated connector proteins) that bind to adhesins. As the MyoA motor is anchored to the IMC, the force generated by the displacement of actin makes the parasite move forward (46). Parasites lacking MyoA are viable, but they show defects in gliding motility as well as in cell invasion and egress, both of which depend on motility (47). In the absence of MyoA, the class XIV Myosin-C and its associated protein GAP80 relocate from the basal complex to the pellicle, where it forms an alternative glideosome complex with GAP50 and GAP40, which accounts for the viability of the MyoA KO (48). Whether MyoC is also replacing MyoA in its role in mitochondrial dynamics is not currently known. Future experiments to address this question would include determining the localization of MyoC upon loss of MyoA and the effect of double MyoA/MyoC knockout on mitochondrial morphodynamics.

Up to now, no role beyond the glideosome has been ascribed to MyoA, particularly in organellar movement. In *Toxoplasma*, the unconventional class XII myosin Myosin-F is involved in organellar transport. Myosin-F is important for apicoplast turnover (20), organelle elongation, and the association of the apicoplast with the centrosomes (19). It is also known that Myosin-F is important for the endomembrane network and secretory system in the parasite (18, 49). Recent studies have shown that Myosin-F is also important for actin dynamics (50). Nonetheless, the involvement of myosin motors in mitochondrial movements is well characterized in other organisms. In yeast, mitochondria-bud-directed movements are driven by Myosin 2 (Myo2). In contrast to MyoA, Myo2 contains a cargo-binding domain. Still, its activity in mitochondrial movements depends on adaptors, such as the Rab-like small GTPase Ypt11 (17) and an OMM-bound protein Mmr1 (33). Mitochondria lacking Myo2 are defective in actin-binding, and mutants of this protein are unable to distribute the organelle during budding, resulting in daughter cells lacking mitochondria. Interestingly, we observed parasites lacking mitochondrial material in the MyoA knockout cell line.

We were able to detect a MyoA-dependent interaction between *Toxoplasma* mitochondria and actin. Furthermore, treatment with either Cytochalasin D or Jasplakinolide affects mitochondrion positioning, supporting a role for actin in *Toxoplasma* mitochondrial morphodynamics. Normally, *Toxoplasma* actin is found primarily in the globular form (51). Polymerization of actin in the parasite is an unusual process due to its structure (52) and kinetics (53), producing unstable F-actin filaments that undergo disassembly and turnover faster than skeletal muscle actin (54). Although unstable, F-actin filaments are important for the development of the parasites in the intracellular niche, forming tubules interconnecting parasites inside of the parasitophorous vacuole and with a possible role in cell-to-cell communication (55). Apicoplast inheritance in *Toxoplasma* is an actin-dependent process. Previous studies have shown that parasites grown in the presence of Cytochalasin D accumulate both apicoplast and apical secretory organelles within the residual body (56). In this work, we are showing for the first time that *Toxoplasma* mitochondrion can interact with actin, confirming the role of the actomyosin system in mitochondrion dynamics. Interestingly, unlike the mitochondria interaction with actin in yeast, which is inhibited by ATP, we observe an interaction between *Toxoplasma* mitochondria and actin regardless of ATP presence or absence. It is still unclear why we haven’t seen an inhibition of actin binding in the presence of ATP. This could be due to differences in the structure of MyoA in comparison to canonical myosins or the particularity of our assays. *Toxoplasma* MyoA has a low affinity for actin in the presence of ATP, and it requires high ATP concentration to reach maximal speed in motility assays (57). Nonetheless, recombinant MyoA can dissociate from actin by the addition of 4 mM ATP (29). Thus, further studies will be needed to determine whether the interaction is indeed ATP-independent.

One important remaining question is the positioning of MyoA in the IMC when interacting with the mitochondrion. Existing data indicates that the glideosome is immobilized on the side of the IMC facing the plasma membrane. Nonetheless, the interaction of the mitochondrion with the IMC involves the side of the IMC facing the cytoplasm. Thus, any involvement of MyoA in the IMC/mitochondrion interaction would require MyoA to be facing the cytoplasmic side of the IMC. Despite the advances in microscopy techniques, such as U-ExM, we still don’t have the level of resolution to determine the exact localization of MyoA within the pellicle. We hypothesize that there might be two populations of MyoA, one located on the cytosolic side, probably interacting with LMF1, and on the plasma membrane side of the IMC, interacting with GAP45. While phosphorylation status is known to influence localization in many proteins, mutations in all phosphorylated amino acids do not alter MyoA’s association with the pellicle (26, 30). Thus, it is plausible that localization is determined by protein-protein interactions.

Phosphorylation of the MyoA motor by CDPK3 is important for motility and egress (26, 58). This post-translational modification (PTM) occurs at specific sites in the N-terminal domain (S20, S21, and S29) and motor domain (S743) (26, 30). Structural studies have shown that the N-terminus phosphorylation promotes protein stabilization and increases actin-based motility (40). On the other hand, phospho-null mutants of these N-terminal sites are defective in calcium-induced egress (26, 30). Our results showed that the N-terminal residue S21 in MyoA is important for correct mitochondrial dynamics. The actin interaction assays showed that mitochondrion extracted from parasites carrying this mutation pulls down more actin, suggesting that this residue is involved in the interaction between mitochondrion and actin. Interestingly, in this mutant, instead of parasites without mitochondrion, we mostly observed parasites containing a branched mitochondrion or even two copies of the organelle. In our model, the S21 phosphorylation might be a fine tune in the differential activity of MyoA. If the residue is not phosphorylated, the interaction with LMF1 occurs. Upon egress signaling, CDPK3 phosphorylates the residue, and the LMF1-MyoA interaction is lost. As the cdpk3-KO cell line (which is not able to phosphorylate S21) does not show the same mitochondrion phenotypes, it is difficult to make conclusions from this set of results.

Based on our result, we proposed that as the mitochondrion branches into the daughter parasites, MyoA interacts with LMF1 to help promote organelle positioning and inheritance. This interaction is dependent on the phosphorylation status of the N-terminal residue S21. As parasites lacking MyoA can still inherit a mitochondrion, it is plausible that other motors might be normally involved in this process or compensate for the loss of MyoA. In mammalian cells, there is a cooperation between microtubules and actin for mitochondrial division and segregation. The kinesin KIF5 is responsible for moving mitochondria along microtubules (59). Overall, there are several putative kinesins and dyneins in the *Toxoplasma* genome (60). Kinesin A is localized to the apical complex, and Kinesin B is localized to 2/3 of the microtubules (24). Both LMF1 Y2H (6) and MyoA proximity biotinylation (this study) suggest an interaction with Kinesin B, but further studies are necessary to characterize the role of this protein in mitochondrion division. As it has been demonstrated, actin dynamics in *Toxoplasma* is a faster process in comparison to mammalian actin (54). It is also plausible that MyoA does not necessarily act as a motor, but as an actin-binding protein, and mitochondrion movements are driven by the actin filaments dynamics.

Our identification of a novel and unique protein-protein interaction that mediates mitochondrial dynamics and inheritance provides the first insights into the active segregation of mitochondrion in *Toxoplasma*. Future studies of other components of this complex, especially those involved in its regulation, will shed light on this important aspect of the biology of Toxoplasma and potentially reveal novel avenues for therapeutic interventions.

## MATERIALS AND METHODS

### Parasite culture and reagents

All the parasite strains were maintained via continuous passage through human foreskin fibroblasts (HFFs) purchased from ATCC (SCRC-1041) and cultured in Dulbecco’s Modified Eagle’s Medium (DMEM) high glucose, supplemented with 10% fetal bovine serum (FBS), 2 mM L-glutamine, and 100 U penicillin/100 mg streptomycin/ml. The cultures were maintained at 37 °C and 5% CO_2_. Parasites used in this include the strain RH (type I) lacking hypoxanthine-xanthine-guanine phosphoribosyl transferase (HPT) and Ku80 (RHDhxgprtDku80) (61), LMF1-HA and LMF1-KO (5) Myosin-A KO (25), FLAG-Myosin-A WT (complement) and FLAG-Myosin-A S21A (26). Cells were inspected for mycoplasma contamination using a Venor^TM^ GeM Mycoplasma Detection kit PCR-based (Sigma, MP0025-1KT).

### Immunofluorescence Assays

For immunofluorescence assays (IFA), infected HFFs were fixed with 4% paraformaldehyde, quenched with 100 mM glycine, blocked, and permeabilized in phosphate-buffered saline (PBS) containing 3% bovine serum albumin (BSA) and 0.25% Triton X (TX-100). Samples were then incubated with primary antibodies diluted in permeabilization solution for 1 hour, washed five times with PBS, and incubated with the respective Alexa Fluor-conjugated antibodies and 5 µg/ml Hoechst 33342 (a nuclear marker; Thermo Fisher Scientific; cat. No. H3570) in PBS for 1 hour. The coverslips were washed five times with PBS. After washes, the coverslips were mounted in Vectashield. Image acquisition and processing were performed using a Nikon Eclipse 80i microscope with NIS-Elements AR 3.0 (objective lens 100×/1.40 Oil Plan Apochromat, *z*-step size of 0.3 µm) or a Zeiss LSM 800 AxioObserver microscope with an AiryScan detector using a ZEN Blue software (version 3.1). The images were acquired using a 63X Plan-Apochromat (NA 1.4) objective lens. All images acquired from this microscope were acquired as *Z*-stacks with an *XY* pixel size of 0.035 µm and a *Z*-step size of 0.15 µm. All images underwent Airyscan processing using ZEN Blue (Version 3.1, Zeiss, Oberkochen, Germany). Images were processed and analyzed using FIJI ImageJ 64 Software. Primary antibodies used for IFA included rabbit anti-HA (C29F4 Cell Signalling, 1:1000 dilution), mouse anti-Myc (9B11 Cell Signaling, 1:1000), rabbit anti-acetyl Tubulin (Lys40) (Millipore ABT241, 1:2000), rat anti-IMC10 (6), mouse anti-F1B ATPase (1:5000, a gift from Dr. Peter Bradley, University of California Los Angeles, CA, USA), rabbit anti-ACP (1:5000,(62)), rabbit anti-TgATOM40 (1:2000, (62)), and mouse anti-SERCA (1:1000,(63)). Secondary antibodies included Alexa Fluor 594- or Alexa Fluor 488-conjugated goat anti-rabbit-IgG and goat anti-mouse-IgG (Invitrogen), all used at 1:1000.

### Biotinylation by Antibody Recognition (BAR)

The BAR protocol was adapted from (8). Freshly lysed parasites were harvested at 1000 x g for 10 minutes, then resuspended in PBS containing 0.1% Tween-20 (PBST) with 4% paraformaldehyde for 20 minutes. The reaction was quenched by the addition of 0.125M glycine in PBS for 5 minutes. Parasites were washed twice with PBST and permeabilized with PBS supplemented with 0.5% triton X-100 for 10 minutes. Parasites were washed in PBST and blocked with 1% BSA in PBST for 2 hours. After this incubation period, samples were incubated with primary antibody resuspended in 1%BSA/PBST overnight at 4 °C. Samples were washed five times with PBST and resuspended in 100 µL containing 1mM biotin phenol (Iris Biotech GmbH) for 10 minutes and then mixed with the previous solution, supplemented with 2mM hydrogen peroxide for five minutes. The reaction was stopped by the addition of 500 mM sodium ascorbate for five minutes. Sampled were washed twice in PBST and resuspended in lysis buffer (PBST with 1.5% SDS and 1% sodium deoxycholate) and heated for one hour at 100 °C. The lysate was clarified by centrifugation (20,000 x g at 4°C) for 10 minutes. The resulting supernatant was incubated with streptavidin beads (Fisher Scientific) overnight at 4°C. The beads were washed twice with PBST, 1M NaCl in PBST two times, and PBS three times. The beads were processed for mass spectrometry.

### Phenotypic characterization of MyoA mutant strains

For the mitochondrial morphology counts, HFFs infected with the inducible knockdown strains were grown for 24 hours. IFA was performed as described above, using mouse anti-F1B-ATPase and rabbit anti-acetylated tubulin. Three mitochondrial morphological categories were quantitated: lasso, sperm-like, and collapsed (3, 5, 6). The other mitochondrial phenotypes observed during the image analysis were counted and categorized as interconnected mitochondrion, amitochondriate parasites, branched mitochondrion, and two individual mitochondria. Experiments were performed in biological triplicates. A total of 150 individual parasites were counted per replicate. For all mitochondrion-related phenotypes, the images comprised 35-40 z-steps, z-stacked and deconvolved.

For the mitochondrial inheritance counts, IFA was performed. Parasites in the late stage of division (two daughter cells completely separated) were counted. The number of parasites with a mitochondrion branch inside was counted. It was considered that a mitochondrion branch was inside when it was possible to observe a mitochondrion that passed the basal complex in a single slice. The images were composed of 35-40 z-steps, z-stacked and deconvolved. A total of 35 parasites were counted per replicate.

### Ultrastructure Expansion Microscopy

Ultrastructure Expansion Microscopy (U-ExM) was performed as described previously (6, 32). Briefly, parasites were seeded in an HFF monolayer grown on glass coverslips in 24-well plates. Primary antibodies used mouse anti-FLAG (9B11 Cell Signalling, 1:500) and rabbit AtpB (AS05 805 Agrisera, 1:2500). Secondary antibodies used in this study were Alexa Fluor NHS 405 NHS-ester, Alexa Fluor 594- or Alexa Fluor 488-conjugated goat anti-rabbit-IgG and goat anti-mouse-IgG (Invitrogen). Secondary antibodies were used at 1:1000, except for the Alexa Fluor NSH-ester 405, which was used at 1:250. Images were acquired from the LSM 800 microscope. Images were AiryScan Processed using the Zen Blue (3.1) software.

### Generation of transgenic parasites

For C-terminal endogenous tagging of LMF1 with a 2x-myc and destabilization-domain (DD), a PacI-flanked fragment of LMF1 (TGGT1_265180) just upstream its stop codon was amplified by PCR and cloned into pLIC-2xmyc-DD-HXGPRT plasmid by In-fusion cloning. XcmI-linearized plasmid (25 ug) was resuspended in P3 buffer (Lonza) and transfected into the FLAG-MyoA WT (complement), S21A, and MyoA-KO parasites using the Lonza Nucleofactor system. Positive transfectants were selected for HXGPRT in a medium containing 50 μg mycophenolic acid per ml and 50 μg xanthine/ml with 100 nM Shld1. Immunofluorescence and western blotting were used to confirm the expression and localization of LMF1.

To generate a fluorescent mitochondrion strain for live imaging, we used a plasmid coding for the red fluorescent protein TdTomato fused to the 55 N-terminal residues of T. gondii heat-shock protein HSP60 (2) with a Chloramphenicol acetyltransferase (CAT) resistance gene (64). Plasmid was NotI-linearized, and 25 μg of plasmid was transfected into the FLAG-MyoA WT (complement), S21A, and MyoA-KO parasites using the Lonza Nucleofactor system. Positive transfectants were selected for CAT in a media with 40 μM chloramphenicol. Positive clones were screened by immunofluorescence to confirm the expression and correct localization of the fused protein.

### Live Imaging

HFFs were split into imaging chambers with a glass bottom. Parasites were allowed to invade host cells 30’ minutes before the experiment. Uninvaded parasites were washed out by three washes in DMEM 10% FBS prewarmed at 37°C. For live cell imaging, growth media was replaced with Fluorobrite DMEM (Thermo Fisher) containing 1% dialyzed FBS, supplemented with 2mM L-glutamine, and 100 U penicillin/100 mg streptomycin/ml, prewarmed to 37 °C. Images were acquired at the Indiana Center for Biological Microscopy using a Keyence BZ-X810 microscope system equipped with a 63X Oil immersion lens and a stage-top environmental incubation system, kept at 37 °C and 5% CO_2_ in a humid atmosphere. Images were acquired every 10 minutes, with a z-step size of 0.15 µm, a total of 13 Z-stacks per image.

### Co-immunoprecipitation assays

To confirm the interaction between LMF1-myc-DD and MyoA, we performed co-immunoprecipitation using the dually tagged cell lines. Intracellular parasites from 2 T75 cultures were syringe-released, by passing through a 21-gauge needle, spun down (1000 x *g* for 10 min at 4°C), washed twice in cold PBS, and resuspended in Pierce co-immunoprecipitation lysis buffer (Thermo Fisher Scientific) with protease/phosphatase inhibitor cocktail (100×, Cell Signaling Technology). After 1 h of lysis at 4°C, the samples were sonicated three times for 20 s each time (20% frequency). After sonication, samples were pelleted (11,000 ***g*** for 15 min), and the supernatant was incubated with anti-myc magnetic beads (Thermo Fisher Scientific). Samples were placed in a rocker for 2.5 hours before beads were washed once with Pierce co-IP lysis buffer and twice with PBS. In the end, the beads and total lysate were resuspended in 2× Laemmli sample buffer (Bio-Rad) supplemented with 5% 2-mercaptoethanol (Sigma-Aldrich) for western blotting.

### Western blotting

Parasite or digitonin fractionated extracts were resuspended in 2X Laemmli sample buffer (Bio-Rad) with 5% 2-mercaptoethanol (Sigma Aldrich). Samples were boiled for 10 minutes at 95°C before separation on a gradient of 4-20% SDS-PAGE gels (Bio-Rad). Samples were then transferred to nitrocellulose membrane using standard methods for either semi-dry or wet transfer (Bio-Rad). Membranes were blocked in 5% milk and probed with primary and secondary antibodies. Primary antibodies used included rabbit anti-HA 1:5,000 (CS29F4 cat. no. 3427S), mouse anti-HA 1:5,000 (6E2 cat. no. 2367S), anti-myc rabbit 1:5,000 (71D10 cat. no. 2278S), anti-myc mouse 1:5,000 (9B11 cat. no. 2276S), anti-FLAG rabbit 1:5,000 (D6W5B cat. no. 14793S), anti-FLAG mouse 1:5,000 (9A3 cat. no. 8146S), and anti-beta actin mouse 1:10,000 (8H10D10 cat. no. 3700S) all from Cell Signaling Technology. Anti-TgATOM40 rabbit G579 (1:5000), anti-TgACP rabbit (1:5000), anti-Tg**-**beta-tubulin (1:5000) (all from Dr. Michael Reese, UT Southwestern), Anti-IMC10 rat 1:5,000 (6), anti-IMC6 rabbit 1:5,000 (from Dr. Peter Bradley, UCLA), anti-eIF2α rabbit 1:10,000 (from Dr. William J. Sullivan, Indiana University School of Medicine), and anti-GFP tag mouse (168AT1211 cat. no. AM1009A, ABGENT) Secondary antibodies included goat anti-rat-IgG horseradish peroxidase (Life), anti-mouse-IgG horseradish peroxidase or goat anti-rabbit-IgG horseradish peroxidase (Sigma-Aldrich) at a dilution of 1:10,000 for one hour (GE Healthcare). Proteins were detected using SuperSignal West Femto substrate (Thermo Fisher) and imaged using the FluorChem R system (Biotechne). Full original western blots are shown in supplemental figure S1.

### Phenotypic characterization of LMF1-2xmyc-DD strains

For the plaque assays, 500 freshly egressed parasites were seeded in a confluent HFF monolayer in 12-well plates, with or without 100 nM Shld1. After five days of incubation, cultures were fixed with methanol for 15 minutes and stained with Crystal Violet. For the Myosin-A KO/LMF1-myc-DD cell line, cultures were stained 10 days post-infection. Plaques were imaged using a Protein Sample imager. The cleared area was calculated using the Colony Area plugin (65) on FIJI software. The experiment was performed in biological triplicates. Doubling assays were performed by immunofluorescence, using IMC3 as a parasite marker. Briefly, freshly egressed ∼20,000 parasites were seeded into HFF monolayers grown in glass coverslips and allowed to invade for 30 minutes. Cultures were washed with DMEM 10% FBS five times to remove the uninvaded parasites and incubated for 48h before fixation with 4% paraformaldehyde in PBS. For each sample, the number of parasites per vacuole was recorded for 50 randomly selected vacuoles.

For the mitochondrial morphology counts, HFFs infected with the inducible knockdown strains were grown in the presence or absence of 100 nM Shld1 for 24 hours. IFA was performed as described below, using anti-mouse anti-rat IMC10 and anti-rabbit ATOM40. Experiments were performed in biological triplicates. For all mitochondrion-related phenotypes, the images comprised 35-40 z-steps, z-stacked and deconvolved.

### Pharmacological disruption of actin and microtubules

To evaluate the effect of actin-disrupting drugs on mitochondrion morphology, parasites were seeded onto HHF monolayers overnight and then treated with 1 µM cytochalasin D or equivalent of DMSO for approximately 16 h before fixation with 4% paraformaldehyde. For Jasplakinolide, parasites grown overnight in HFF monolayers were treated with 1 µM Jasplakinolide for 2 hours before fixation.

### Subcellular fractionation and mitochondrion enrichment of *Toxoplasma*

The mitochondrion enrichment protocol was performed as previously described (34) with some modifications. Briefly, T175 flasks were infected with parasites from the strains SOD2-GFP/IMC1-TdT (35), LMF1-HA and LMF1-KO (5), MyoA KO (25), and MyoA WT (Complement) and MyoA S21A (26). After 36 hours of infection, intracellular parasites were syringe-released from host cells using a 21-gauge needle and harvested by washing twice (1,000 g for 10 minutes at 4°C) with a Sorbitol-Tris-EDTA (SoTE, 0.6M, 20 mM, and 2 mM respectively) buffer, pH=8. Parasites were counted and resuspended in SoTE to a final concentration of ∼1.5×10^8^ parasites. Part of the sample (50 μL) was removed for the total lysate control. Then, cells were fractionated by the addition of SoTE buffer containing 0.1% (final concentration 0.05%) or 0.2% (final concentration 0.1%) of digitonin with Protease Inhibitors (ThermoFisher). Samples were mixed by pipetting and incubated on ice for 5 minutes. Cell suspensions were separated by centrifugation (8000 g for 5 minutes at 4°C), and the pellet was separated from the supernatant. A fraction of the supernatant was also reserved for downstream assays. Pellets were resuspended in SoTE containing Protease Inhibitors cocktail and kept on ice for further assays.

### Actin interaction assays

Actin co-sedimentation assays were performed as previously described (13) but with the following modifications. Enriched mitochondrion samples were washed once in the actin dilution buffer (5mM Tris-HCl pH=8, 0.2 mM CaCl_2_, and 2mM MgCl2) (8,000 g for 5 minutes at 4°C) and resuspended in the actin reaction buffer (15 mM Tris-HCl pH=7.5, 25 mM KCl, 10mM MGCl_2_ and 0.1mM EGTA). Pre-formed actin filaments (Cytoskeleton, Inc.) were handled according to the manufacturing instructions. Samples were incubated with 100 ug/mL F-actin for 30 minutes in the presence or absence of ATP (10 mM). The reaction was stopped in a buffer containing 25% sucrose and 20 mM HEPES-KOH, pH=8.0. Samples were centrifuged at 12,000 x *g* for 5 minutes at 4°C, and all the supernatant was removed. Samples were resuspended in 2X Laemmli Buffer with 5% 2-mercaptoethanol and boiled for 10 minutes at 95°C.

For the visualization of the interaction by immunofluorescence, the mitochondrion vesicles were pre-incubated with anti-rabbit ATOM40 antibody (1:2000), and actin filaments were pre-stained with ActinGreen 488 reagent (phalloidin, ThermoFisher, #A12379). After the staining, the interaction assays were performed as described above. As a secondary antibody, Alexa Fluor 594-conjugated goat anti-rabbit-IgG was used. Image acquisition and processing were performed using a Nikon Eclipse 80i microscope with NIS-Elements AR 3.0 (objective lens 100×/1.40 Oil Plan Apochromat, z-step size of 0.3 µm).

## ACKNOWLEDGMENTS

The mass spectrometry work performed in this work was done by the Indiana University School of Medicine Center for Proteome Analysis. The live imaging in this article was obtained using the facilities of the Indiana Center for Biological Microscopy. This research was supported by the National Institutes of Health grants R01AI149766, R01DK124067, and R21AI164619 to G.A. The live imaging work was funded, in part, with support from the Indiana Clinical and Translational Sciences Institute funded, in part by Grant Number UL1TR002529 from the National Institutes of Health, National Center for Advancing Translational Sciences, Clinical and Translational Sciences Award to R.O.O.S.

---

**Figure S1.**
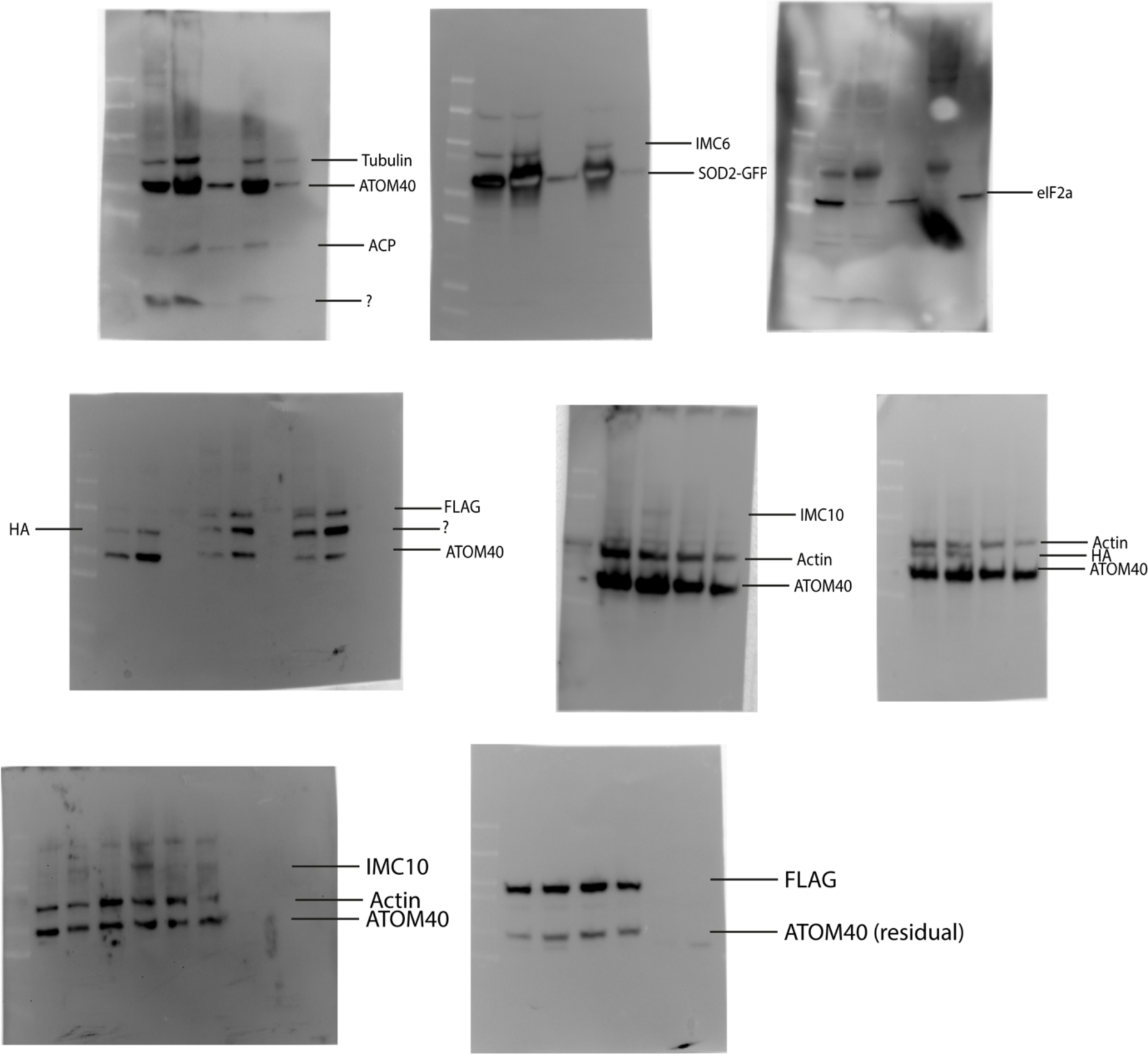
– Raw images of western blots. Western blots of Figure 6 (A) and Figure 10 (B).

**Figure S2.**
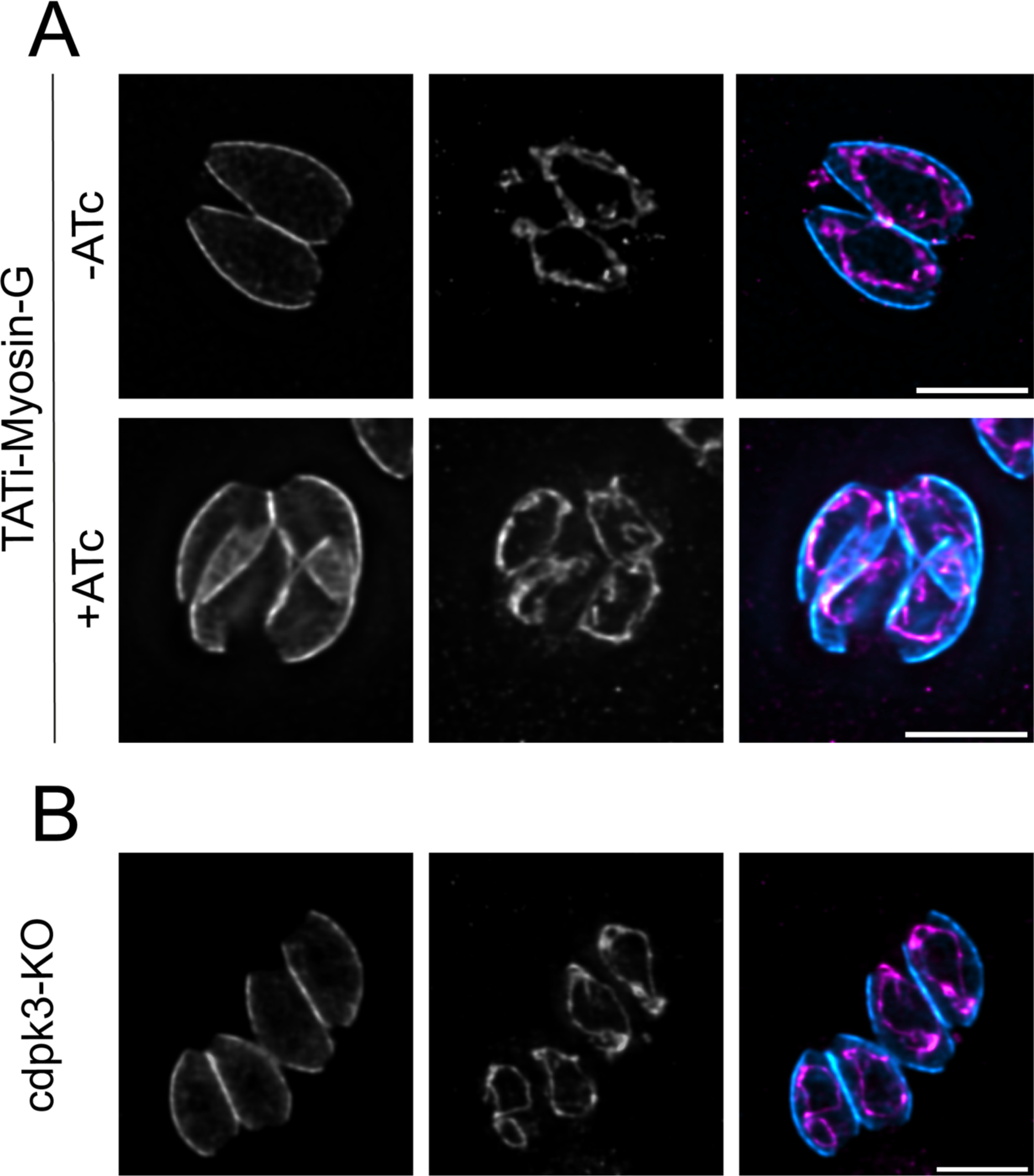
– Depletion of Myosin-G and CDPK3 does not affect mitochondrion morphology in intracellular parasites. Intracellular parasites of the TATi-Myosin-G (A) and cdpk3-KO (B) were grown in fibroblasts for 16 hours, then stained for IMC10 (cyan) and F_1_B-ATPase, a mitochondrial marker (magenta). To induce the knockdown of Myosin-G, parasites were kept in the presence of anhydrotetracycline (ATc, 0.5 µg/ml) for 16h. Scale bar: 5 μm.

**Figure S3.**
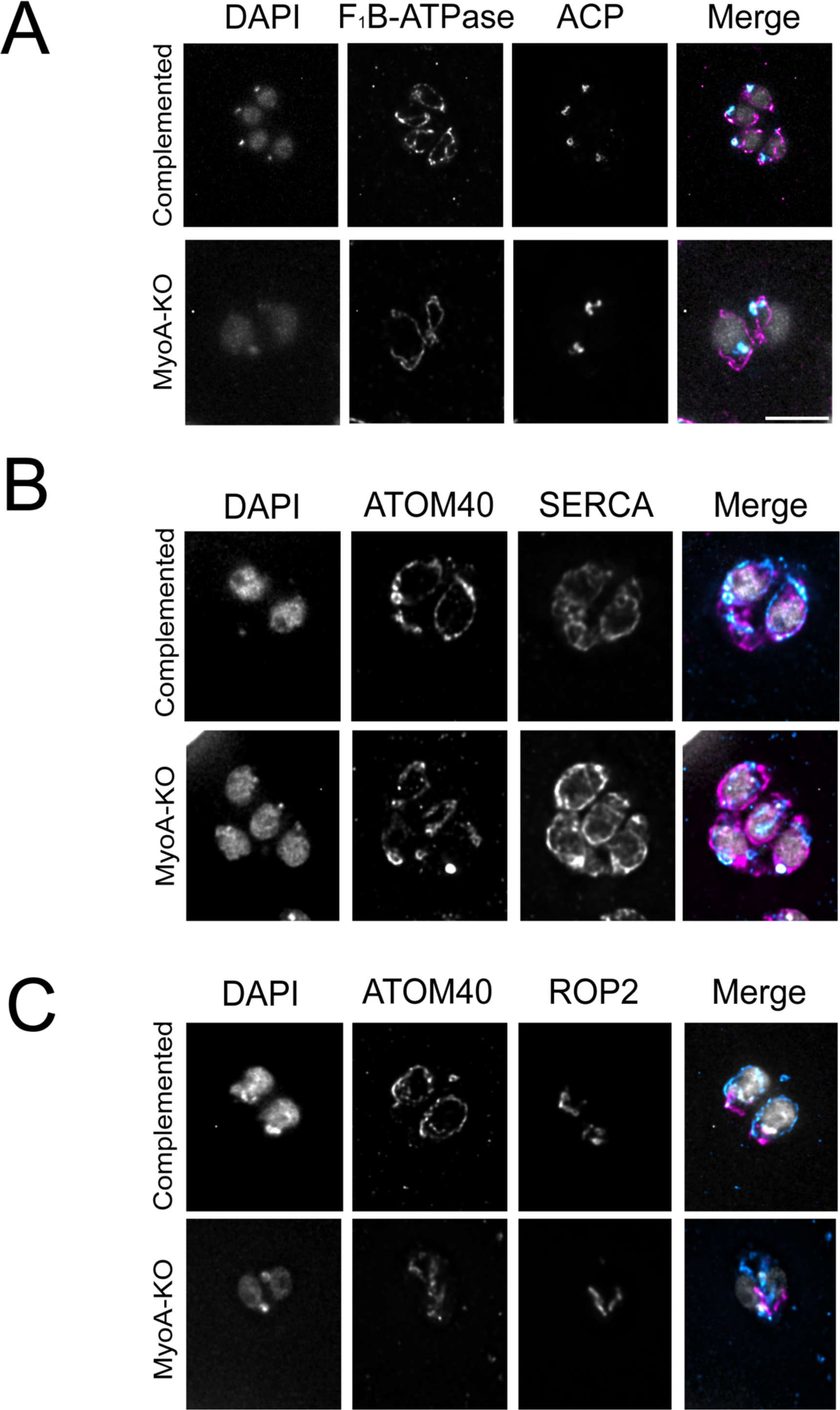
Depletion of Myosin-A does not affect apicoplast, endoplasmic reticulum, or rhoptries morphology in intracellular parasites. Intracellular parasites of the complement and myosin-A KO strains were grown in fibroblasts for 16 hours, then stained with anti-ACP to detect the apicoplast (A), anti-SERCA to detect the ER (B), or with anti-ROP2 to detect the rhoptriesROP2 (rhoptries). Either F_1_B-ATPase (A) or ATOM40 (B and C) were used as a mitochondrial marker. Scale bar: 5 μm.

**Figure S4.**
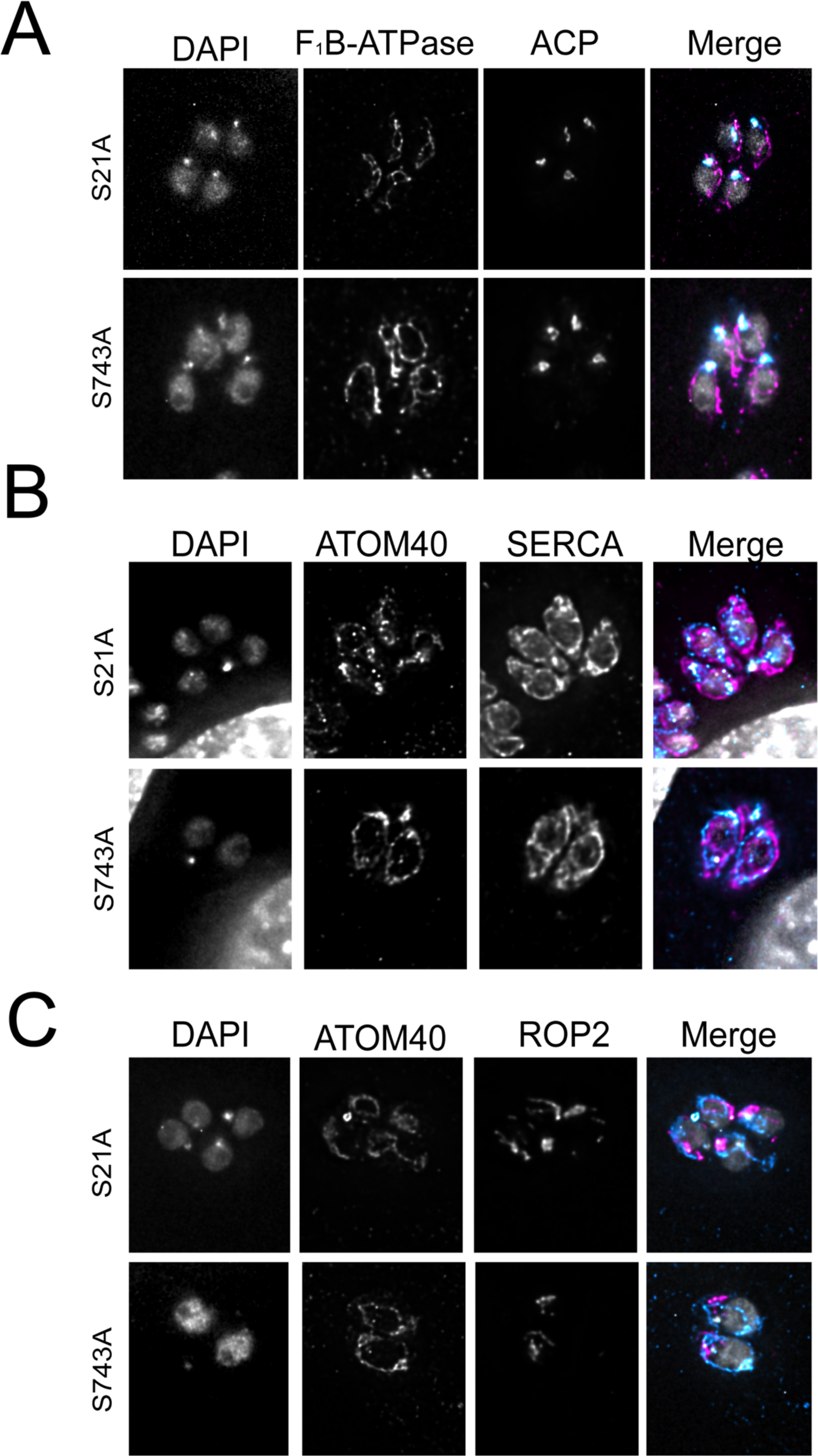
N- or C-terminal mutation of Myosin-A does not affect apicoplast, endoplasmic reticulum, or rhoptries morphology in intracellular parasites. Intracellular parasites expressing FLAG-Myosin-A S21A and FLAG-Myosin-A S743A were grown in fibroblasts for 16 hours.Cultures were then stained with anti-ACP to detect the apicoplast (A), anti-SERCA to detect the ER (B), or with anti-ROP2 to detect the rhoptriesROP2 (rhoptries). Either F_1_B-ATPase (A) or ATOM40 (B and C) were used as a mitochondrial marker. Scale bar: 5 μm.

**Figure S5.**
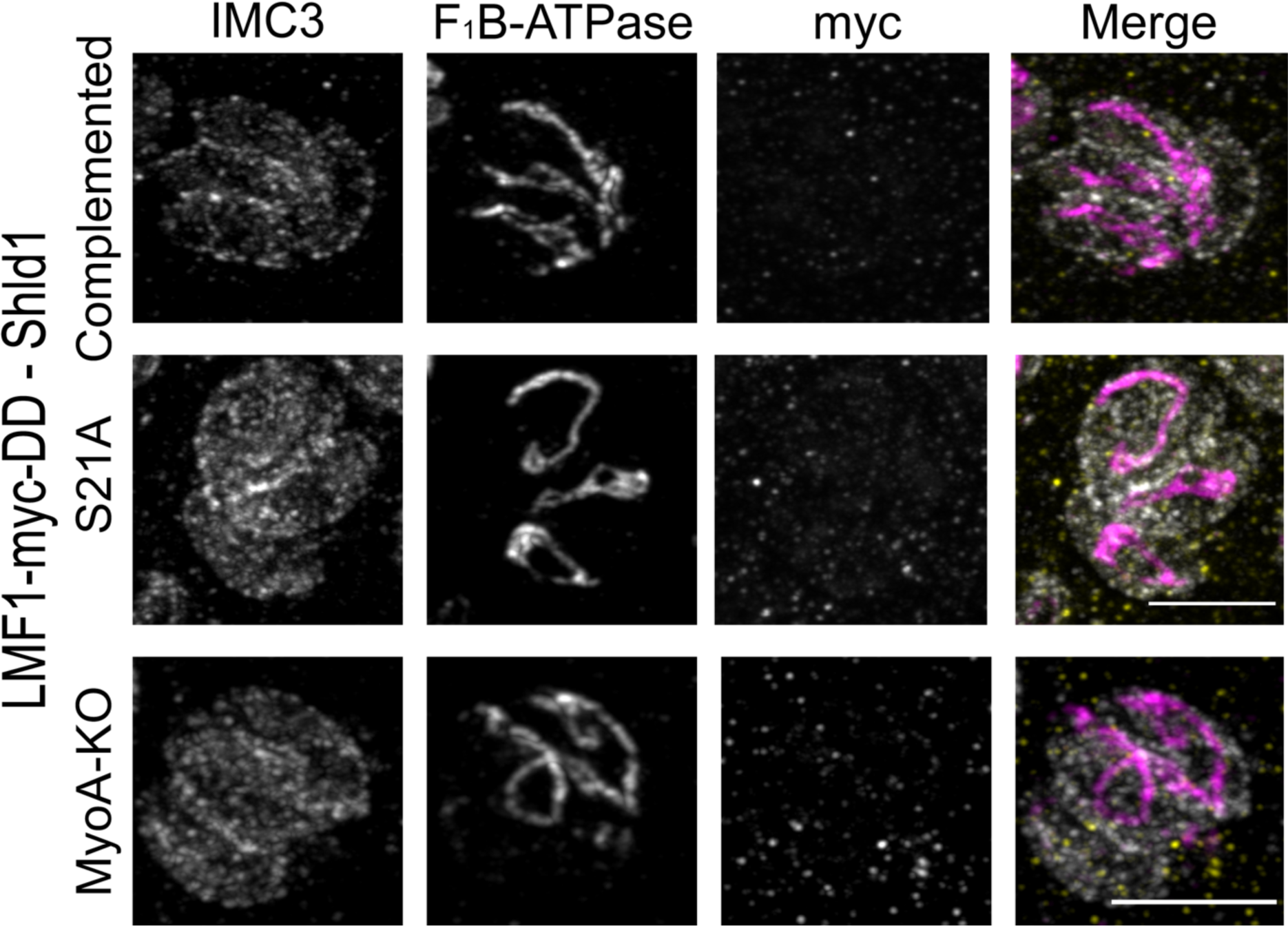
Establishment of parasite strains with conditional expression of LMF1. Representative IFA of MyoA complemented, MyoA S21A, or MyoA KO parasites expressing LMF1-myc-DD showing expression LMF1(Myc) in the absence of Shld1. Parasites were stained for IMC3 (gray), F_1_B-ATPase (magenta), and myc (yellow). Scale bar: 5μm.

**Figure S6.**
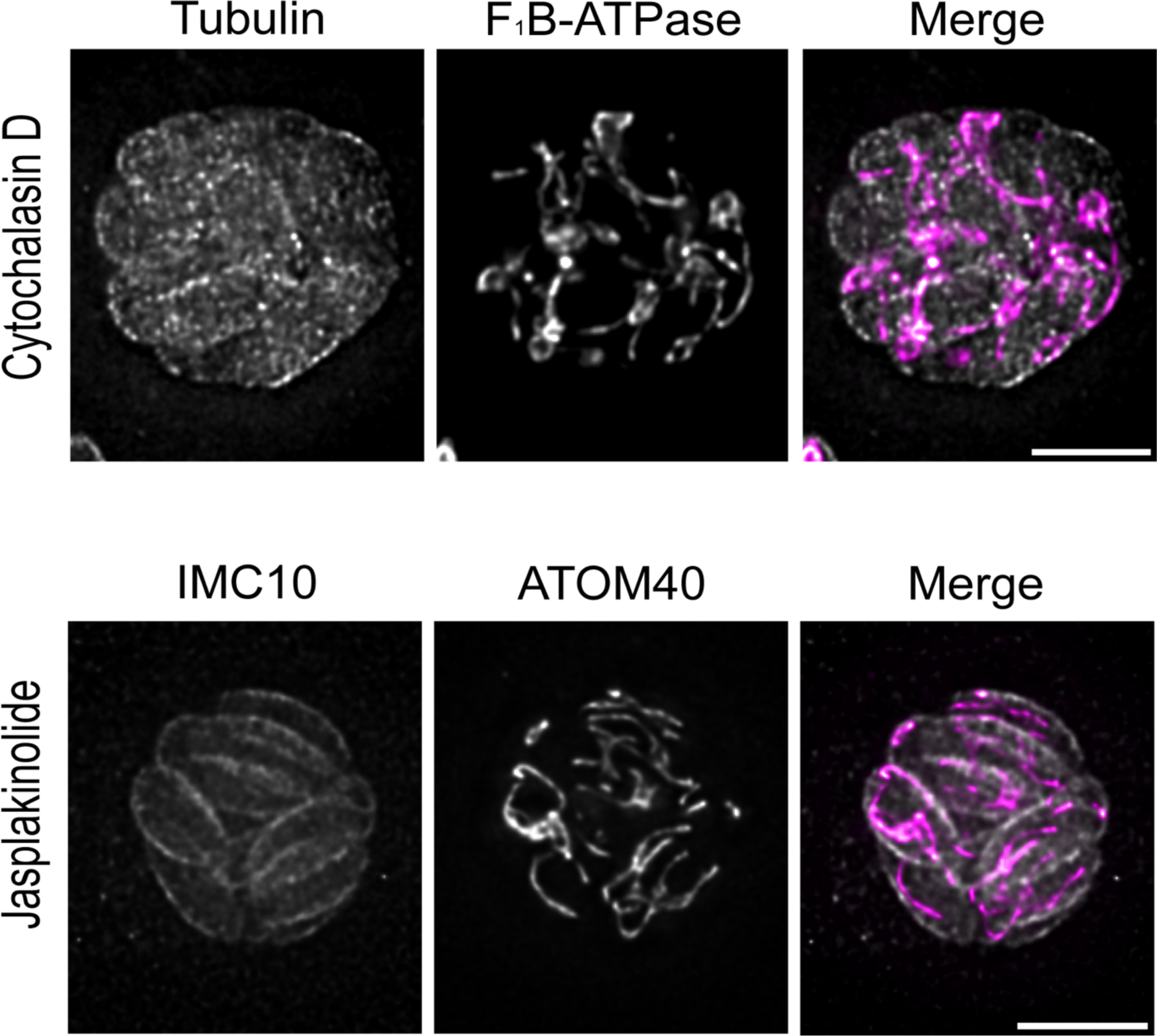
Actin is critical for mitochondrion morphology. IFA of large parasite vacuoles grown in the presence of 1 mM of Cytochalasin D (A) or 1 mM Jasplakinolide (B). Parasites were stained for Tubulin (gray) and F_1_B-ATPase (magenta) or IMC10 (gray) and ATOM40 (magenta). Scale bar: 5μm.

**Figure S7.**
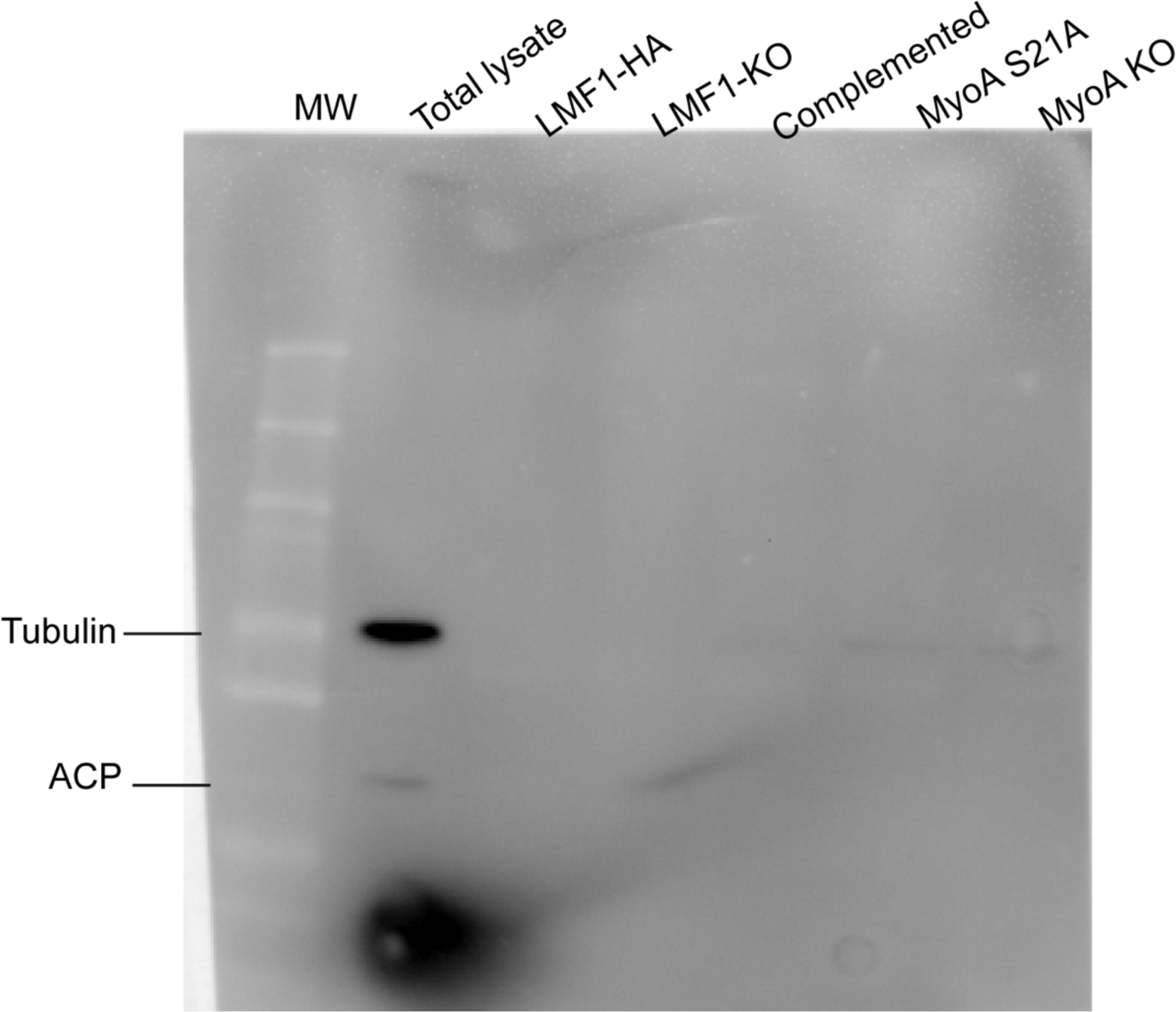
Enriched mitochondrion fractions do not carry microtubules and apicoplasts. Membranes of post-cushion enriched mitochondrion vesicles were probed for the presence of microtubules (anti-Tubulin) or apicoplast (anti-ACP).

